# Whole-body integration of gene expression and single-cell morphology

**DOI:** 10.1101/2020.02.26.961037

**Authors:** Hernando M. Vergara, Constantin Pape, Kimberly I. Meechan, Valentyna Zinchenko, Christel Genoud, Adrian A. Wanner, Benjamin Titze, Rachel M. Templin, Paola Y. Bertucci, Oleg Simakov, Pedro Machado, Emily L. Savage, Yannick Schwab, Rainer W. Friedrich, Anna Kreshuk, Christian Tischer, Detlev Arendt

**Affiliations:** Developmental Biology Unit, European Molecular Biology Laboratory (EMBL), Meyerhofstrasse 1, Heidelberg, 69117, Germany; Sainsbury Wellcome Centre for Neural Circuits and Behaviour, Howland Street 25, London, W1T 4JG, UK; Cell Biology and Biophysics Unit, European Molecular Biology Laboratory (EMBL), Meyerhofstrasse 1, Heidelberg, 69117, Germany; Heidelberg Collaboratory for Image Processing, Institut für Wissenschaftliches Rechnen, Ruprecht Karls Universität Heidelberg, Heidelberg; Friedrich Miescher Institute for Biomedical Research, Maulbeerstrasse 66, Basel, 4058, Switzerland; Princeton Neuroscience Institute, Princeton University, Washington Rd, Princeton, NJ, 08544, USA; Department of Neuroscience and Developmental Biology, University of Vienna, Althanstr. 14, Vienna, 1090, Austria; Centre for Ultrastructural Imaging, King’s College London, New Hunt’s House Guy’s Campus, London, SE1 1UL, UK; Cell Biology and Biophysics Unit / Electron Microscopy Core Facility, European Molecular Biology Laboratory (EMBL), Meyerhofstrasse 1, Heidelberg, 69117, Germany; Centre for Bioimage Analysis, Core Facilities, European Molecular Biology Laboratory (EMBL), Meyerhofstrasse 1, Heidelberg, 69117, Germany

**Author notes:** These authors contributed equally (Hernando M. Vergara, Constantin Pape, Kimberly I. Meechan, Valentyna Zinchenko, Christel Genoud, Adrian A. Wanner).

**Keywords:** volume electron microscopy, image registration, automatic segmentation, gene expression atlas, *Platynereis dumerilii*, tissue biology, ultrastructure, multimodal data integration, machine learning

## Abstract

Animal bodies are composed of hundreds of cell types that differ in location, morphology, cytoarchitecture, and physiology. This is reflected by cell type-specific transcription factors and downstream effector genes implementing functional specialisation. Here, we establish and explore the link between cell type-specific gene expression and subcellular morphology for the entire body of the marine annelid *Platynereis dumerilii*. For this, we registered a whole-body cellular expression atlas to a high-resolution electron microscopy dataset, automatically segmented all cell somata and nuclei, and clustered the cells according to gene expression or morphological parameters. We show that collective gene expression most efficiently identifies spatially coherent groups of cells that match anatomical boundaries, which indicates that combinations of regionally expressed transcription factors specify tissue identity. We provide an integrated browser as a Fiji plugin to readily explore, analyse and visualise multimodal datasets with remote on-demand access to all available datasets.

## Introduction

Cells are the basic units of life. In multicellular organisms, distinct sets of genes are expressed in different cells, producing their individual cellular traits that we call cell types (Arendt et al., 2016). Deciphering how the genotype is decoded into a multicellular phenotype is therefore critical to understand the development, structure and functioning of an entire body. Towards this goal, we need to establish the link between expression profiles and cellular morphologies. This requires techniques that permit the integration of genetic and phenotypic information for all cells of the body. On one hand, volume electron microscopy (EM) techniques have recently produced 3D ultrastructural cellular information in unprecedented coherency and detail (volume scanning EM reviewed by (Titze and Genoud, 2016), serial section transmission EM of a full *Platynereis* larva by (Randel et al., 2014)). On the other hand, spatial single-cell -omics techniques have revolutionised expression profiling in the past years (Lein et al., 2017). The aim of this study is to integrate both in a multimodal cellular atlas for an entire animal body, which has not yet been achieved.

Integration of EM with functional data has been successfully accomplished through correlative light electron microscopy - as exemplified for the mouse cortex (Bock et al., 2011; Holler-Rickauer et al., 2019; Lee et al., 2016) and retina (Bae et al., 2018; Briggman et al., 2011), or the zebrafish olfactory bulb (Wanner and Friedrich, 2020). However, correlative light electron microscopy has not yet been extended at larger scale to also include transcriptional information at the single cell level (but see (Jahn et al., 2016)). In reverse, spatial transcriptomics protocols do not yet sufficiently preserve samples for ultrastructural analysis by EM. Therefore, the integration of gene expression and cellular ultrastructure remains a big challenge, especially at the full organism scale.

Here, we introduce a pipeline to achieve this for the entire body of a 6 days post fertilisation (dpf) young worm of the marine annelid *Platynereis dumerilii*. At this stage, *Platynereis* already exhibits a rich and differentiated set of cell types, which is comparable to that of many bilaterians, including vertebrates. However, since each cell type comprises few cells, the overall number of cells remains small, which entails a much smaller body size than that of vertebrates or insects of similar complexity. This goes in concert with a considerable stereotypy of *Platynereis* development and differentiation: the developmental lineage is invariant (Vopalensky et al., 2019), and differentiated larvae and young worms resemble each other down to the cellular detail (Asadulina et al., 2012; Randel et al., 2015; Tomer et al., 2010; Vergara et al., 2017). We first acquired a serial block-face scanning electron microscopy (SBEM) volume for a whole animal, and introduced a novel semi-automated multi-scale approach to produce a complete cellularly segmented volume of the animal. We then expanded an existing gene expression atlas for the 6 dpf stage (Vergara et al., 2017), and registered it to the segmented EM volume.

For the first time, this enabled us to assign gene expression information to segmented cells throughout an entire animal. Building on this, we examined the correlation of gene expression and cellular morphology for diverse tissue types. We demonstrate that an unbiased gene expression clustering defines groups of neurons that form coherent tissues separated by morphological boundaries. Focusing on the animal head, we identified molecular-anatomical domains that correspond to brain ganglia. Importantly, in order to coherently integrate, efficiently explore, and easily analyse the data derived from multiple modalities (EM data, single cell segmentation, gene expression, custom annotations, cell type definition, etc), we implemented an open-source Fiji-based plugin that we call MultiModal Browser (MMB). We enable the community to browse and mine the resources described here for *Platynereis dumerilii*, by providing an online browsing tool, based on the MMB, that we refer to as PlatyBrowser. We expect our novel workflow to be transferable to EM volumes of other animals showing some degree of developmental stereotypy, and the tools presented here to generate a multimodal catalogue of cell types across organisms.

## Results

### A whole-body serial block-face scanning electron microscopy dataset

An EM image stack of a complete young worm of *Platynereis dumerilii*, late nectochaete stage (6 dpf), was collected by SBEM at a pixel size (x/y) of 10 nm and 25 nm section thickness (z). Each planar image was composed of multiple tiles, the number and distribution of which was adapted to the surface of the animal cross-section. The automated image acquisition process was controlled by the open-source software SBEMimage (Titze et al., 2018) and took seven weeks. The resulting stack contains 11,416 planar images with >200,000 tiles, and has a total size of 2.5 TB. This dataset enabled detailed analyses of both overall anatomy and fine ultrastructural details throughout the whole organism (**Figure 1**), allowing for the visualisation of characteristic subcellular features of different organs and cell types (**Figure 1B-E, Supplementary Figure 1**). To demonstrate the quality of the EM data, a selection of cells and their features are highlighted below.

**Figure 1:**
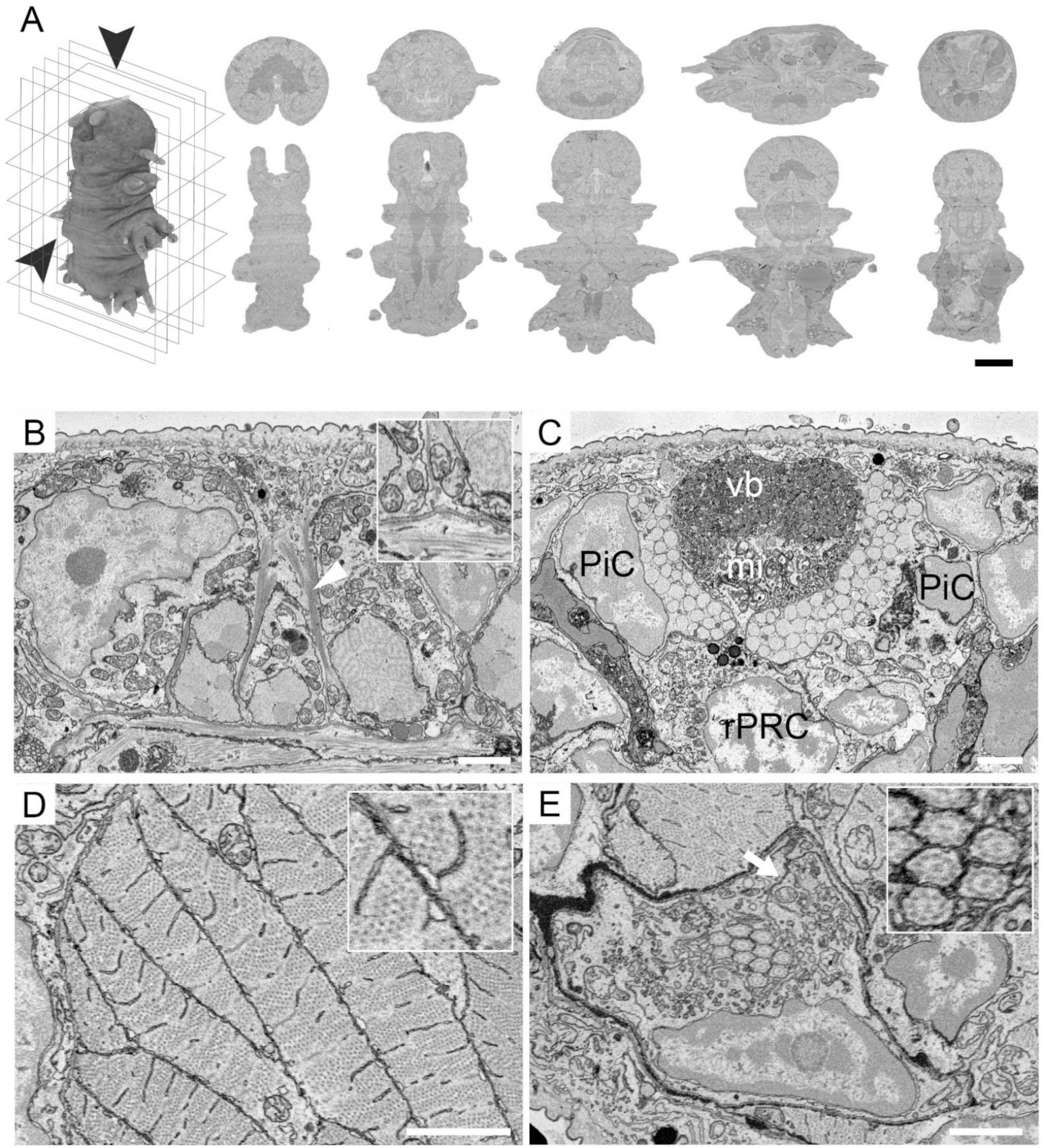
Ultrastructure of a whole *Platynereis* by serial block-face scanning electron microscopy. **A**: The 3D SBEM dataset can be observed in multiple orientations, either along the acquisition plane (transversal plane, top row) or as orthogonal projections (horizontal plane, bottom row; scale bar: 50 µm). **B-E**: fine ultrastructure is revealed when exploring the datasets at native resolution (10 nm pixel-size x/y; scale bars: 2 µm). **B**: epithelial cell, interfacing the cuticle and the underlying muscle. Bundles of cytoskeletal filaments (arrowhead) form part of the attachment complex (inset). **C**: the adult eye forms a pigment cup composed of pigment cells (PiC) and rhabdomeric photoreceptors (rPRC). The photoreceptors extend a distal segment made by microvillar projections (mi) for light detection. In the centre of the pigment cup is the vitreous body (vb). **D**: longitudinal muscle fibres are cut transversally displaying cross-sections of the sarcomere as well as of the sarcoplasmic reticulum that contacts the plasma membrane (inset). **E**: cross section of the distal part of the nephridia, highlighting the autocell junction (arrow) which forms a lumen. The lumen houses a bundle of motile cilia (identified by the 9+2 microtubule arrangement, inset), which are contributed by each cell of the nephridium, noted by the presence of basal bodies. Panels B to E are snapshots selected from the full volume and can be retrieved directly from the PlatyBrowser “Bookmark” function.

Epithelial cells surround the body of *Platynereis* supporting the cuticle. These cells display attachment complexes made of bundles of cytoskeletal fibres, likely intermediate filaments, which span from the apical to the basolateral side of the cell, thus connecting the cuticule to the underlying muscle layer (**Figure 1B**). At the epithelial to muscle junctions, anchoring points can be identified (**Figure 1B inset**), where plasma membrane deformation of both the epidermal and muscle cells reflects the mechanical forces exerted at this interface. Intermediate filaments are also visible in the ciliated support cells (**Supplementary Figure 1A**). These cells in the head, posterior to the adult eyes, are part of the sensory nuchal organ (Chartier et al., 2018; Schmidtberg and Dorresteijn, 2010). They show an enrichment of mitochondria at their apical side, and similar to epithelial cells, they form junctions with the underlying muscles (**Supplementary Figure 1A inset**).

Other organs which can be identified based on morphologically distinct cells are the adult eyes (**Figure 1C**). At 6 dpf there are 4 adult eyes, grouped as pairs located dorsally on each side of the head, immediately underneath the cuticle. These eyes are composed of pigment cells and rhabdomeric photoreceptors, which together form a pigment-cup type eye (Rhode, 1992). The characteristic microvilli of rhabdomeric photoreceptors, which increase the surface area of these cells for light detection, are distinguishable within each eye, as are the sub microvillar cisternae of the endoplasmic reticulum and the vitreous body of the eye.

Moving internally to the mesoderm, the longitudinal muscles are clearly identifiable. They are ventrally located and sectioned transversally in the imaging plane (**Figure 1D**). Individual myosin filaments are visible but actin filaments cannot be resolved because their size is below the image resolution. These muscles lack the t-tubule system present in vertebrates and instead have an extended sarcoplasmic reticulum, as described by (Rosenbluth, 1968). The sarcoplasmic reticulum appears tubular when spanning the myocyte and forms a disc parallel to the sarcoplasmic membrane (**Figure 1D inset**).

The larval nephridium (also called ciliated protonephridia (Hasse et al., 2010)) is found adjacent to the longitudinal muscles, located laterally between the 2nd and 3rd larval segments (**Figure 1E, Figure 2D, Supplementary Figure 1B**). It is composed of 7 cells on each side of the body which connect to the coelomic cavity. Each cell forms a lumen, which is closed off via auto-cell junctions (**Figure 1E**), and into which they protrude several motile cilia (identifiable by the 9 + 2 microtubules, **Figure 1E inset**). Proximally, the first cell projects several cilia into the coelomic cavity, and other cilia into a bundle that travels distally in the nephridial lumen to the next cell of the nephridia. Each subsequent cell contributes new cilia, as proven by the presence of basal bodies. The terminal cell of each nephridium forms an external pore for excretion, the nephridiopore, which is surrounded by microvilli. Each nephridial cell contains multiple vesicles mediating various forms of cell transport (Bartolomaeus and Quast, 2005). Sites of endocytosis between the cells of the nephridia and the coelomic cavity can be identified (**Supplementary Figure 1B inset**).

**Figure 2:**
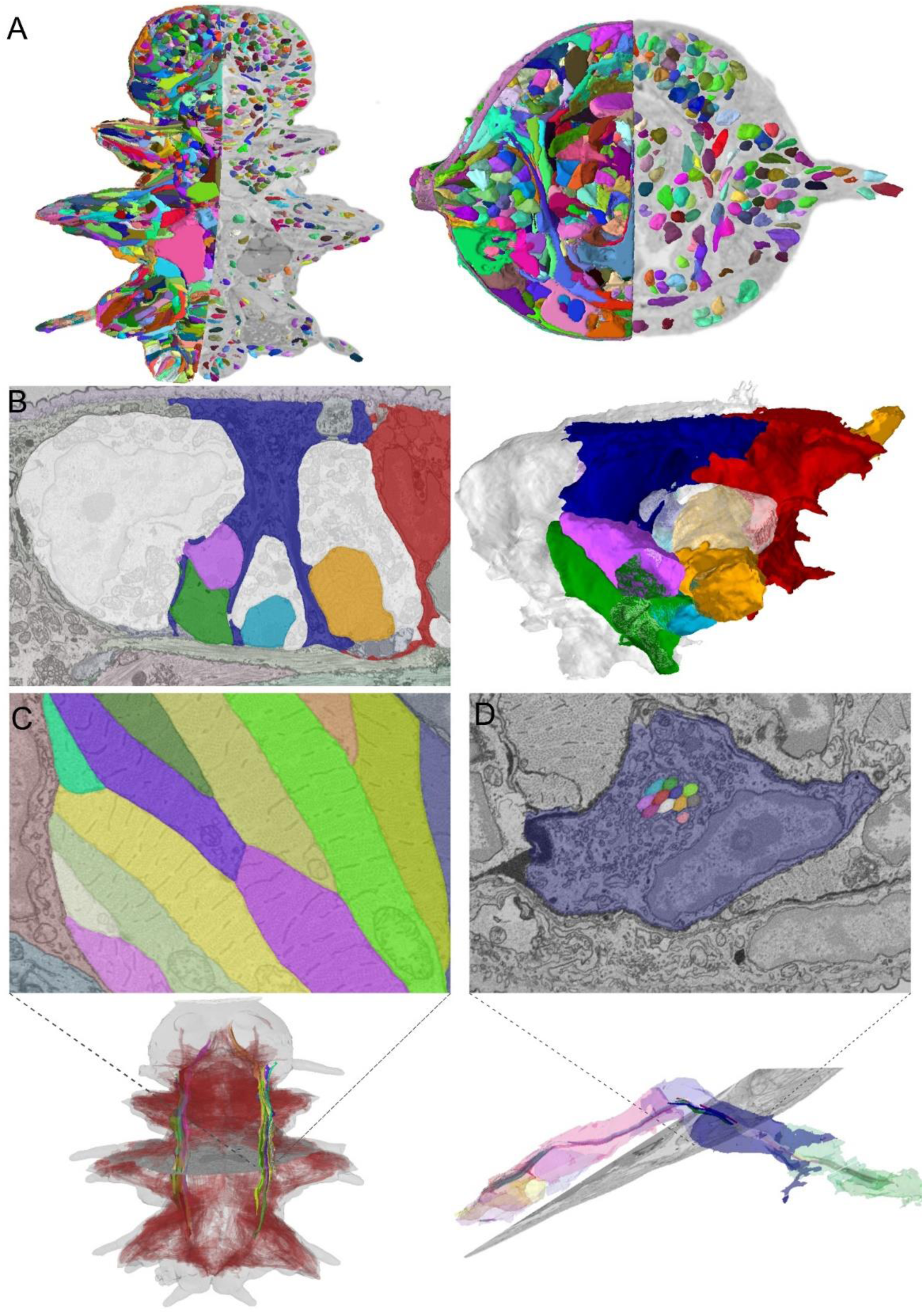
Segmentation of nuclei, cells, tissues and body parts. **A**. Cells and nuclei are segmented as 3D objects in the whole EM volume. Horizontal section (left) and transversal section (right) with 3D renderings of all cells (left half) and nuclei (right half). The cellular segmentation yields accurate 3D reconstructions for the different cell types in the animal. **B**. Intertwining epithelial cells are reconstructed, see coloured segments overlayed with EM on the left and 3D rendering on the right. **C**. Long stretching muscles are segmented correctly, see overlay with EM in the top image. This bundle of muscles is highlighted in the bottom rendering, with the corresponding bundle on the other side coloured less brightly and all other reconstructed muscles shown as brown renderings in the background. **D**. We studied the nephridia in more detail and also reconstructed all cilia that contribute to their central cilia bundle. Seven cells contribute to this bundle, see the top image for an intersection of one of these cells and the bundle and the bottom image for a 3D rendering of the seven cells and cilia where each cilium is coloured the same as the cell it is attached to. For panels B, C and D bookmarks are available in the PlatyBrowser.

Similar ultrastructural analysis can be performed on each cell throughout the full dataset, illustrating the resolution power of SBEM imaging on the full 6 dpf *Platynereis*. The entire dataset is made available for download via an open source data repository (EMPIAR 10365) and it is browsable via the PlatyBrowser described in detail below.

### Segmentation of nuclei, cells, tissues, and body parts

Having acquired a complete specimen at ultrastructural resolution, we aimed to segment every cell in the body, as well as subcellular structures. Nuclei were segmented by an extension of the mutex watershed algorithm from (Wolf et al., 2018), based on pixel affinity predictions of a 3D U-Net (Çiçek et al., 2016). For the cells, we have extended the Lifted Multicut-based approach, introduced for neuron segmentation in EM by (Beier et al., 2017), to segment a much larger variety of cells found in the whole-animal volume. Large-scale brain segmentation pipelines commonly perform the segmentation of cells in a purely bottom-up manner, based on the prediction of cellular membranes provided by a convoluted neural network (CNN) (Funke et al., 2019; Januszewski et al., 2018; Lee et al., 2017). We found this approach not to be sufficiently robust to variability in cell membrane appearance across the whole animal. The effect of this variability can be mitigated by drastically increasing the CNN training data volume - a requirement that is hard to satisfy as the training data has to be obtained by dense manual segmentation in 3D. Instead, we exploited the automatically segmented nuclei to introduce top-down constraints as additional edges in the Lifted Multicut framework following the approach of (Pape et al., 2019) and thus induced additional cell boundaries derived from the nucleus segmentation. In addition to the nuclei-based constraints, we take advantage of the multi-scale nature of our data by segmenting the tissues which are clearly discernible at lower resolutions: muscle, neuropil, glands, the gut and the flat surface epithelium. Tissue boundaries are then also added to strengthen the boundaries in the cell segmentation problem.

The automatic segmentation algorithm produced 11,402 cells with nuclei (**Figure 2A**). These were scored based on their morphology, and the 154 most implausible cells - which turned out to be falsely merged with their neighbours - were corrected in a semi-automatic manner. In addition, we used the Paintera software (Hanslovsky et al., 2020) for very fine-grained corrections of several cells of interest such as the nephridia described below. The segmentations were validated against 8 manually annotated slices (4 transversal, 4 horizontal) distributed throughout the dataset. Here, we found a 99.0% agreement with the automatic nuclear segmentation and a 90.3% agreement with the cellular segmentation (see Methods and **Supplementary Figure 2A**). Figure 2 gives a few examples of the achieved segmentation quality for epidermal cells (B), muscles (C) and nephridia (D). We measured nuclei sizes to range from 33.6 to 147.5 cubic microns, and cell sizes from 59.8 to 1224.6 cubic microns. Note that neurites in the neuropil have not been segmented, as they are not sufficiently preserved in the EM volume for automated segmentation.

In addition to the tissue segmentation (see **Figure 2C** for the entire musculature segmentation), we subdivided the body (and cells) into head, the first body segment (cryptic segment), the ventral nerve cord, the unsegmented posterior part of the body (pygidium), the lateral ectoderm, the foregut and the midgut (see **Supplementary Table 1**). Since at 6 dpf the head is already morphologically highly diverse (Chartier et al., 2018), we also segmented the head into morphologically discernible subunits or ganglia. These include the paired palpae, antennae, cirri, and adult eyes representing sensory organs, and central complex as well as mushroom bodies as brain parts (**Supplementary Figure 2D**). The detail in the 3D ultrastructure volume and the cell segmentation provide an unprecedented resolution and framework to do this anatomical classification in a complete and unbiased manner (as compared to previous efforts recognising brain anatomy mostly based on immunohistochemistry (Chartier et al., 2018; Tomer et al., 2010)).

To evaluate the quality of cellular segmentation, we focused on the larval nephridia (**Figure 2D**). Previous transmission EM analysis had shown that the larval nephridia are embedded between the body surface muscles and oblique muscles, and that the tubule wall is formed from single cells wrapping around a tight lumen with six cilia constantly present (Hasse et al., 2010). However, transmission EM on single sections could not resolve how many cells contribute to this structure and how many distinct cilia they protrude into the lumen. This task is now possible with our SBEM resource: we segmented all nephridial cilia as proof of principle that comprehensive ultrastructure segmentation is possible from the SBEM resource. The two nephridia stereotypically comprise 7 cells per side, and each cell contributes several cilia to the continuous central bundle. The bundles are made up of 85 and 78 cilia on the left and right sides of the body, respectively. Furthermore, we observed a similar distribution of cilia per cell for both sides and found that for a given cross-section of the lumen cilia belong almost exclusively to one cell (**Supplementary Figure 2C**).

Finally, the pronounced contrast in subnuclear structure also allowed a semantic segmentation of chromatin using the nuclear segmentation as a mask (**Supplementary Figure 2E**). Many nuclei showed a meandering pattern of light and dark subregions, corresponding to classical euchromatin and heterochromatin plus nucleolus.

### Morphological clustering of segmented cells

The whole-body cellular-scale segmentation allowed us to quantitatively describe and compare the morphology of all cells. To do this, we defined and measured 140 morphological descriptors for all segmented cells and nuclei, including the chromatin distribution inside the nucleus (see **Supplementary Table 2**). These descriptors include features of the size, shape, intensity and texture of segmented objects, such that every cell was represented as a vector of size 140 describing its position in morphology space. We then clustered all cells using a graph-based approach (see methods), and visualised with UMAP (Uniform Manifold Approximation and Projection for Dimension Reduction) (McInnes et al., 2018). This analysis demonstrated a broad separation into morphologically discernible tissues. We found these to be largely congruent with the segmented tissues such as neural tissue, musculature, epidermis and midgut tissue (**Supplementary Figure 2F**). Notably, we observed some further subdivisions of neurons based on morphology, which is remarkable in the absence of neuropil segmentation (and thus, any information about projection). For example, we found cluster 1 is almost entirely confined to the head, with an enrichment in one of the segmented head ganglia (ganglia 1). Conversely, clusters 2, 5 and 7 were more mixed containing neurons from a number of regions including the head, foregut and ventral nerve cord. The morphological differences between the perikarya of cluster 1 and the other neural clusters were mostly texture features of the chromatin and the intensity of the nucleus (see **Supplementary Table 2**).

We next explored the power of morphometry with regard to the most stringent, smallest grouping possible – that of bilateral cell pairs. In the *Platynereis* 6 dpf young worm, the pronounced stereotypy of hundreds of distinct cell types in an animal with relatively few cells leads to the presence of bilateral pairs of cells across the entire animal (Vergara et al., 2017). To detect the bilateral partner of a given cell, we ranked all other cells in the organism in terms of their distance in morphology space (from first nearest neighbour, to most distant neighbour). We then identified potential bilateral partner cells by their mirror-image position in the body (see methods for details). Strikingly, based on all morphology features combined, 14% of cells found a potential bilateral partner within the 5 nearest neighbours (20% within 10 and 28% within 20 nearest neighbours; **Supplementary Figure 2B**). This is remarkable given that for each cell there are more than 10,000 partners to choose from in this analysis. While all features combined performed the best, followed by nuclear features (shape, intensity, texture and chromatin distribution), we noted that the chromatin features alone (intensity and texture) still found bilateral pairs with surprising efficiency (e.g., 7% of cells finding a partner within the 5 nearest neighbours, 12% within 10 and 18% within 20). Since bilateral pairs of cells in the 6 dpf *Platynereis* are also identified via identical expression profiles (Vergara et al., 2017), this would suggest some correlation between chromatin morphology and gene expression profile.

### A comprehensive cellular gene expression atlas

The stereotypic pattern of cellular division and differentiation, both temporal and spatial, of *Platynereis* development (Vopalensky et al., 2019), allows for whole-body generation of gene expression atlases with cellular resolution based on image registration and Profiling by Signal Probability mapping (ProSPr) (Achim et al., 2018; Asadulina et al., 2012; Vergara et al., 2017). To obtain comprehensive expression information for all cells in the body at 6 dpf, we introduced new genes into this resource representing areas with poor expression coverage in previous versions of the atlas, such as the digestive system. For example, we selected 37 genes that we found highly enriched in a foregut cDNA library (see methods) and that proved to be differentially expressed in the foregut. We also introduced additional genes with preferential nervous system expression, with special focus on neural differentiation genes (supplementary information). As a result, the 6 dpf ProSPr gene expression atlas (referred to as ProSPr atlas) now contains information for 201 genes (including 78 transcription factors, 56 neural effector genes, and 53 other differentiation genes) and 4 EdU proliferation stainings, which provide an overall good representation of expression profiles throughout the body (**Figure 3A** and **Supplementary Figure 3**). We provide gene expression maps in BioStudies (accession number S-BIAD14).

**Figure 3:**
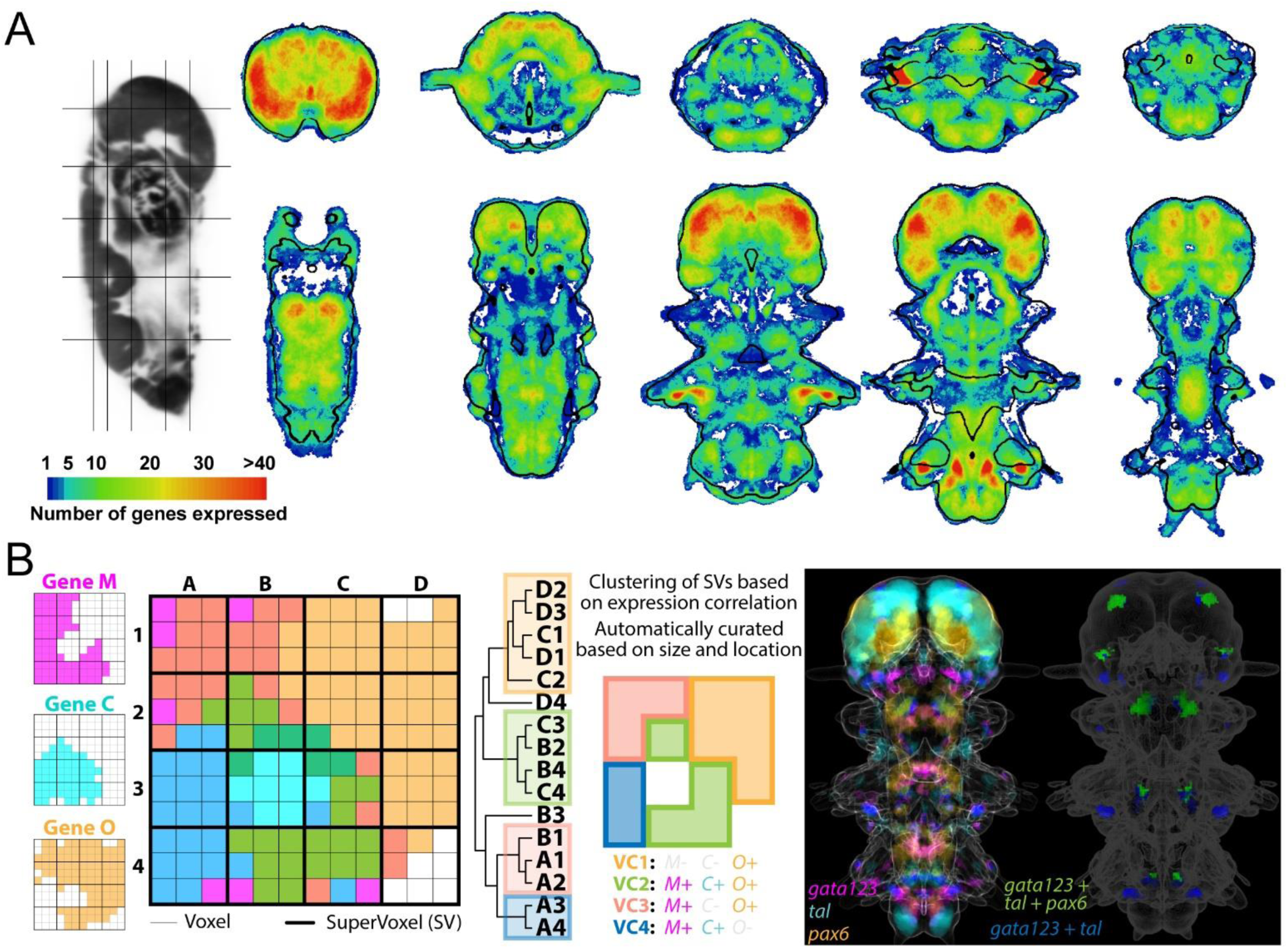
A comprehensive cellular gene expression atlas. **A**: Genetic coverage throughout the animal. Transversal and horizontal sections colour-coded based on the amount of expression information, in gene number, for every pixel. Black contours outline the DAPI-based reference, thresholded for illustration purposes. **B**: Generation of Virtual Cells. On the left, hypothetical spatial expression of three different genes in a 12 × 12 voxel array. In the matrices, thin lines demarcate the voxels and dark ones supervoxels. Voxel colours indicate all possible combinations of expressions (e.g. gene C + gene M in dark blue). The hierarchical tree illustrates the process of clustering supervoxels based on expression information, which renders groups of supervoxels called Virtual Cells (VCs), that are then automatically curated based on size and spatial location. The VCs are visualised spatially on the matrix next to the tree with their expression information. On the right, illustration of this procedure with real data. Three genes are shown on a projection image of the full dataset using similar colouring for the co-expression as in the example on the left. Next to it, colouring of VCs showing a specific expression pattern (2 examples). Note that each of these groups is composed of many VCs but all have the same colouring for illustration purposes.

To identify cell representatives within the ProSPr atlas, we subdivided the expression space into units of homogenous gene expression (Vergara et al., 2017). For this, the expression space was subdivided into SuperVoxels (SVs), cubes of 27 isotropic voxels of 0.55 µm side (each SV represents a cube of 1.65 µm edge length and 4.5 µm^3^ volume). This ensures a representation of each cell by a minimum of 12 SVs (see methods). We then calculated the gene expression for each SV (**Figure 3B**), and clustered the SVs based solely on gene expression, selecting groups of SVs that matched the average size of 1 to 6 cells, as some cell types are segmentally repeated (Vergara et al., 2017). We found the majority of clusters comprising spatially confined groups of SVs with bilateral symmetry, which is a good indication of the high quality of the data. In order to extract individual cell representatives from these clusters, each cluster was then split into spatially connected objects, which are further automatically curated based on size and bilateral symmetry (see Methods). This process generated 12,393 separated 3D entities (called Virtual Cells), a number very similar to that of EM-segmented cells (11,402).

### Registration of SBEM stack with the gene expression atlas

We next set out to integrate the SBEM volume dataset with the ProSPr atlas. As the typical variation between individuals in the position of individual cells is less than one cell diameter (<4.7µm) (Vergara et al., 2017), we reasoned that registration should be possible at close to cellular resolution. To avoid compromising the appearance of ultrastructural morphology, we computed a transformation of the lower resolution ProSPr atlas onto the higher resolution SBEM volume.

We used the software package elastix (Klein et al., 2010; Shamonin et al., 2013) to compute a multi-step registration of the average DAPI signal (representing nuclei of 153 specimens from the ProSPr atlas) onto the binary mask of the segmented nuclei of the EM individual. To assess the accuracy of this registration, we manually identified 43 corresponding landmarks between the datasets and determined their spatial discrepancies in the registered datasets (supplemental information). First, we computed a rigid similarity transformation, only allowing for rotation, translation, and a uniform scaling factor. Comparing the datasets matched in this manner already highlighted the high stereotypy of 6 dpf *Platynereis* larvae as the same micrometre-sized features were visible in the aligned datasets (**Figure 4A**). However, it also revealed expected systematic differences, possibly due to differences in sample preparation. This rigid transformation resulted in a median discrepancy between landmarks of 10.09 µm (more than twice the average cell diameter). To improve the registration, we next allowed for local deformation using a sequence of BSpline (Rueckert et al., 1999) transformations with a final grid spacing of 5.5 µm. This reduced the median discrepancy of the landmarks to 2.99 µm (less than one cell diameter). Applying BSpline transformation to the ProSPr atlas thus allows for the overlay of expression information onto ultrastructure data at cellular resolution.

**Figure 4:**
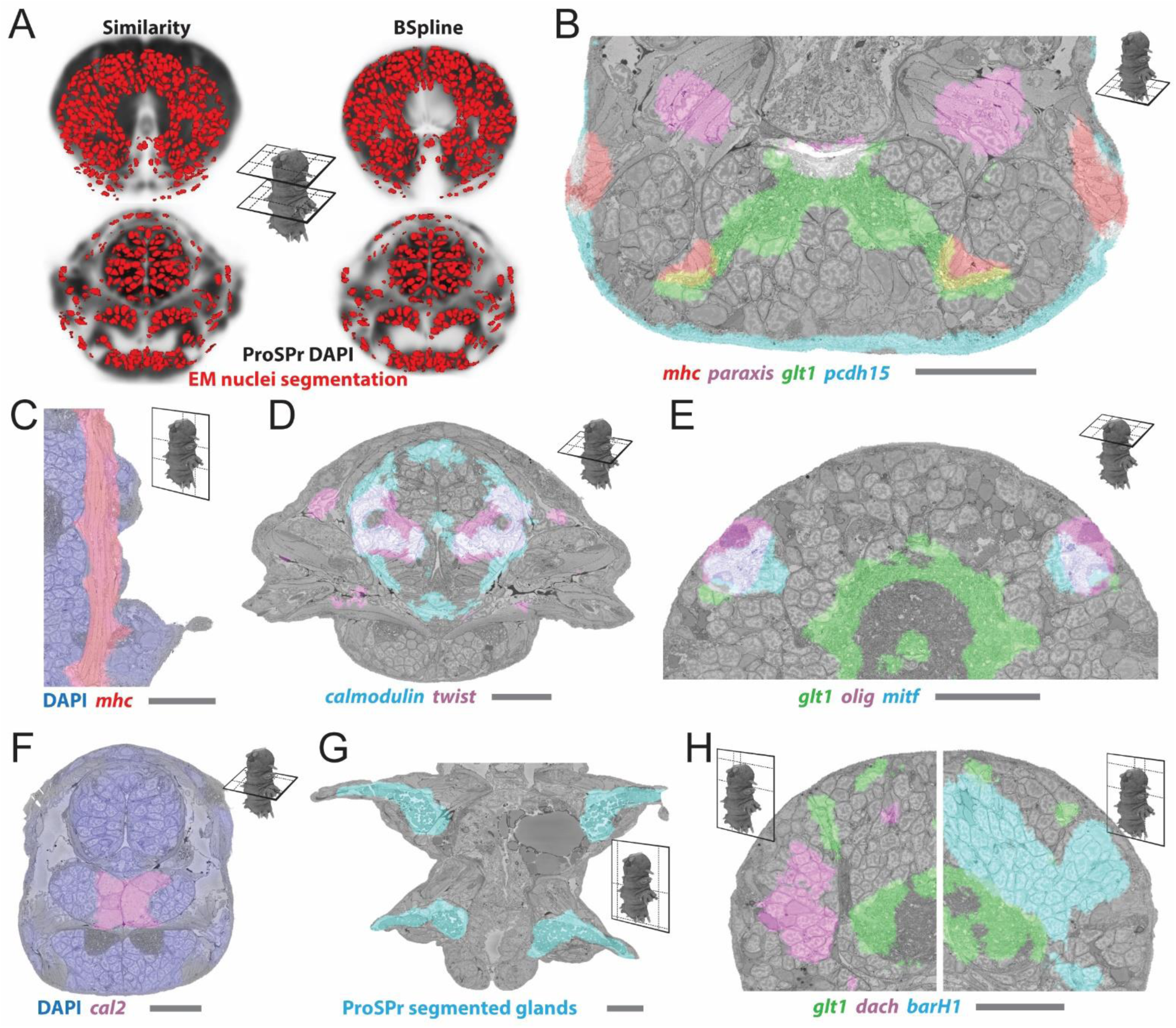
Registration of ProSPr atlas with the EM dataset. **A**: Comparison between similarity and BSpline transform, illustrated with two transversal slices. ProSPr DAPI reference (grey tones) with overlaid segmented EM nuclei (red). Top: cross-section through the head region; bottom: cross section through the foregut region. Cross-section locations are illustrated in the centre of the image. **B - H**: Diverse examples of overlay of different genetic markers with structurally-distinct animal regions evident from the EM. For each panel, the genes are indicated below. DAPI refers to the ProSPr DAPI reference signal. ProSPr segmented glands (G) were extracted from the ProSPr dataset alone using the autofluorescence of the animal. Scale bar is 25µm in all images. The location of the cross-section is illustrated as in A. For panels B-H bookmarks are available in the PlatyBrowser.

An overlay of genetic markers with corresponding tissues confirms the registration accuracy. For example, muscle gene expression is confined to cells with striated and smooth muscle ultrastructure across the body. Expression of the *myosin heavy chain (mhc)* gene overlays the longitudinal and other muscles (**Figure 4B,C**); the *paraxis* transcription factor labels the oblique parapodial muscles and the axochord (**Figure 4B**); and the *twist* transcription factor localises to a subset of muscle cells in the stomodeum (**Figure 4D**). We also find the average distribution of some neural transcripts, such as the glutamate transporter *glt1* (**Figure 4B,E,H**) and the nicotinic acetylcholine receptor (*nAchR*), accurately overlapping the neuropil, indicating that these transcripts are transported into neurites. Furthermore, epithelial genes such as the cadherin *pcdh15* (**Figure 4B**), the metabotropic glutamate receptor *grm7* and the serine protease Neurotrypsin *ntrps*, overlap as expected with the flat epithelial cells lining the body (Achim et al., 2018). Another example shows the cathepsin like L-protease *cal2* localising to oral gland cells (**Figure 4F**) that show similar ultrastructure to the salivary glands in other annelids (Walz et al., 1988). Other expression patterns appear to obey tissue boundaries within an organ, such as the calcium-binding *calmodulin* in the stomodeum (**Figure 4D**) and the neuronal transcription factors *dachshund (dash)* and *barH1* in nervous tissue (**Figure 4H**). Structures like the trunk glands clearly overlap between the two datasets (**Figure 4G**). Finally, we used the adult eyes for registration validation, with their complex cellular morphology comprising rhabdomeric photoreceptor cells and pigment cells (**Figure 1C**) (Rhode, 1992). Our registration reveals two major patterns of averaged gene expression in the morphologically identifiable eyes, one representing pigment plus photoreceptor cells (*mitf*), and the other one representing photoreceptor cells alone (*olig*) (**Figure 4E**).

### Correlation of gene expression with morphologically defined tissues

The unique combination of cellularly-resolved gene expression and available ultrastructure allowed us to address the interplay between differential gene expression and morphology across the cells of an entire body. To do so, an expression value was calculated for every gene in every segmented cell as the fraction of the cell’s volume overlapping the registered gene expression volume (referred to as ‘overlap assignment’). This value, running from 0 to 1, can be interpreted as the probability of expression for a particular gene in a given segmented cell, with higher overlap values denoting a higher confidence that a particular gene is expressed. We then used a graph-based clustering approach (with visualisation by UMAP) that resulted in 15 clusters (**Figure 5A**). To understand how these clusters relate to the animal’s anatomy, we mapped gross body parts (head, cryptic segment, ventral nerve cord, pygidium, lateral ectoderm, foregut and midgut) onto the same UMAP and found a strict correlation with gross morphology, with six genetic clusters spanning the head and two the ventral nerve cord (**Figure 5B**). We also mapped the gene expression-based clusters onto the 3D EM volume, and found segmented cells that belong to the same genetic cluster occupy spatially coherent territories in the organism with a clear correspondence of genetic and tissue boundaries (**Figure 5C**). In the trunk, these transcriptionally defined domains correspond to ventral nerve cord and peripheral ganglia, musculature and gland fields. In the head, we observed a more refined subdivision into central and peripheral domains.

**Figure 5.**
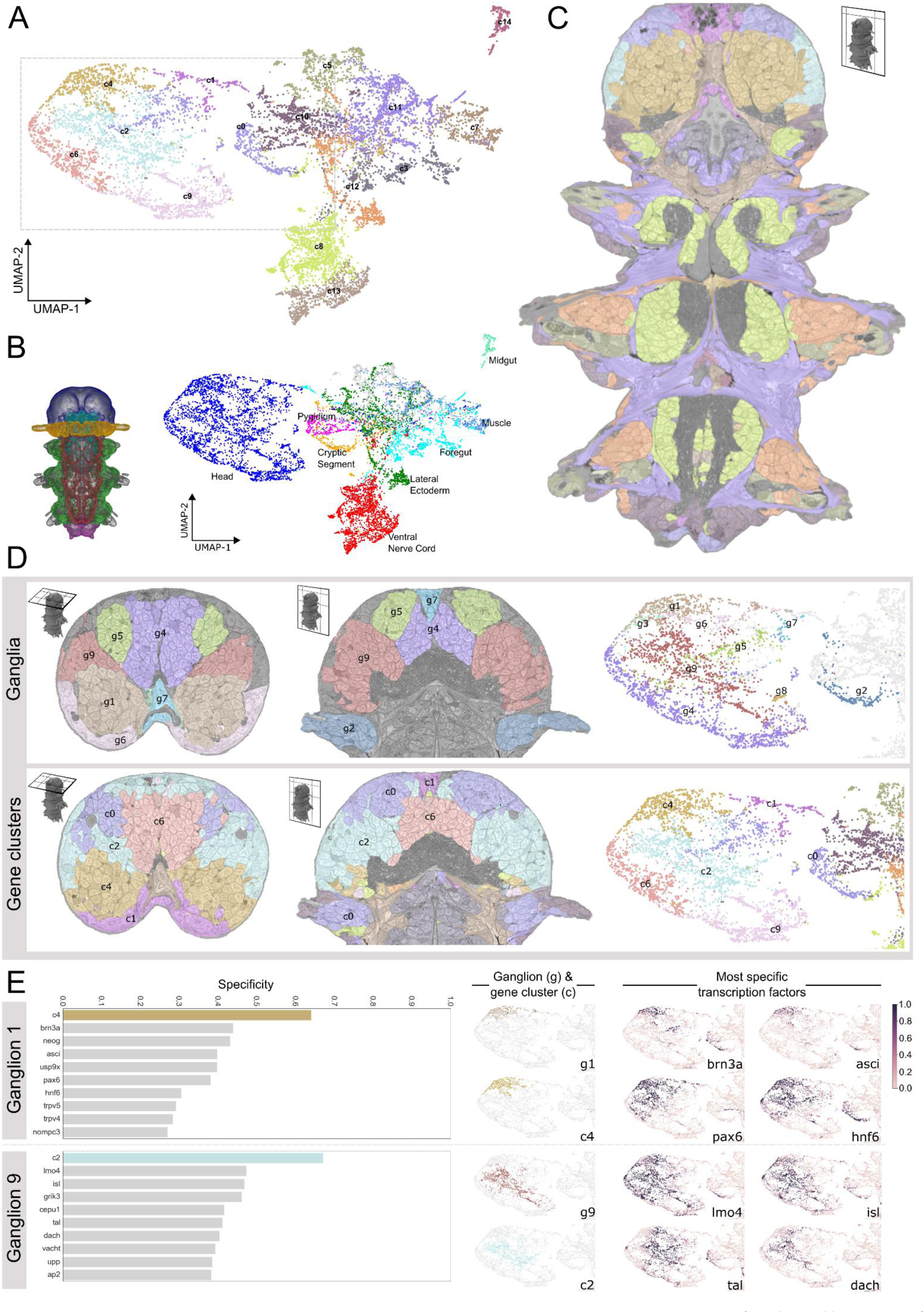
Correlation of gene expression with morphologically defined tissues. **A.** UMAP of all cells based on gene expression data of all 201 genes in the atlas. Points are coloured by membership to different gene expression clusters (c0 – c14). The grey rectangle shows the part of the UMAP that is repeated in panels D and E. **B.** Left – diagram of a *Platynereis*, with broad anatomical regions in different colours (head, cryptic segment, ventral nerve cord, lateral ectoderm, foregut, pygidium). Right - UMAP from panel A, with points coloured by region / tissue type. **C.** Gene clusters overlain on a section of the EM dataset. **D.** Comparison of morphologically defined ganglia (top row), and genetically defined clusters (bottom row). Left / middle column - sections of the EM dataset coloured by membership to different ganglia (top) or gene expression clusters (bottom). Right column – zoom of the head region of the UMAP from A coloured by ganglia, g1-g9 (top) or gene clusters, c0-c14 (bottom). **E.** Comparison of specificity of gene clusters and individual genes for two head ganglia (g1 – top row, g9 – bottom row). Left - graphs of the top 10 scoring genes (grey bars) or gene clusters (coloured bars) by F1 specificity score (see methods). Right – zooms of the head region of the UMAP from A coloured by ganglia (g), top scoring gene cluster (c), or gene expression overlap value (0-1) for top scoring transcription factors. Note: here we show only the head region of the UMAPs, for easier comparison, but some genes and gene clusters have expression domains outside of the head which contribute to their lower specificity scores. See **Supplementary Figure 5** for specificity of the remaining ganglia. For panel C, and the EM overlays of panel D, bookmarks are available in the PlatyBrowser.

For a more detailed analysis of how gene expression relates to tissue boundaries, we focused on the Platynereis head (**Figure 5D**) – the region with the highest gene density in the expression atlas (**Figure 3A**). The *Platynereis* brain is subdivided into morphologically discernible ganglia that we segmented from the EM volume (g1 to g9, **Figure 5D** upper row, **Supplementary Figure 2D** and see above). Returning to our gene-based clustering (**Figure 5D** lower row), we noted that the 6 head clusters (c0, c1, c2, c4, c6, c9) and their respective transcriptional domains in the head showed a marked spatial correlation to these ganglia when overlaid on the EM volume, and also when compared on the UMAP plots. To quantify this correspondence, we calculated a ‘specificity’ score for each genetic cluster for every ganglion (**Figure 5E** and **Supplementary Figure 5**). A highly spatially correlated cluster will confine most of its extent to a given ganglion, and cover most of the area of that ganglion. Hence, our specificity score was calculated as an F1 score (see methods) combining the percent of cluster cells that belong to a given ganglion, with the percent of that ganglion covered by cluster cells. To benchmark these scores, we also calculated specificity scores for all cluster-specific genes (**Figure 5E** and **Supplementary Figure 5**). Remarkably, for almost all ganglia the specificity values for the individual genes were considerably lower than the specificity values for the gene clusters (exceptions were ganglia 3 and 8 that represent very small ganglia). Genes with relatively high specificity values often encoded transcription factors such as *brn3, ascl*, or *pax6* (**Figure 5E**). Characterising these factors further we noted that expression occurs in coherent and overlapping domains, but covers several of the cluster-defined territories. These findings indicated that the *Platynereis* 6dpf head is subdivided into transcriptional domains that are defined by combined expression of several transcription factors (but not by any single factor). These domains largely correspond to morphologically distinct brain ganglia.

### Assigning virtual cells to segmented cells

*Platynereis* body is composed of thousands of cell pairs with homogenous gene expression (see above). These pairs show bilateral symmetry and likely represent cell types (Vergara et al., 2017). Similarly, our nearest neighbour analysis of segmented cells based on morphological descriptors often reveals pairs of cells in bilaterally symmetrical arrangement (**Supplementary Figure 2B**). With the aim of identifying cell types in our dataset more comprehensively, we set out to correlate gene expression specifically with individual cells. Using overlap assignment, this requires individual decisions on whether the measured overlap between each cell and gene expression area (or expression probability) suffices to consider the gene expressed in that cell (**Figure 6A**). At the level of the individual cell, these decisions are necessarily hampered by the biological variability of the EM specimen and by random and systematic deformations that occur during fixation and sample preparation for EM (**Supplementary Figure 6A**). As a result, expression of the same gene may reach the chosen threshold on one side of the animal but remain at subthreshold levels on the other side, which leads to asymmetric assignments (illustrated for *msx* and *patched* in **Figure 6A** and for *wnt5* and *lhx6* in **Figure 6B**).

**Figure 6.**
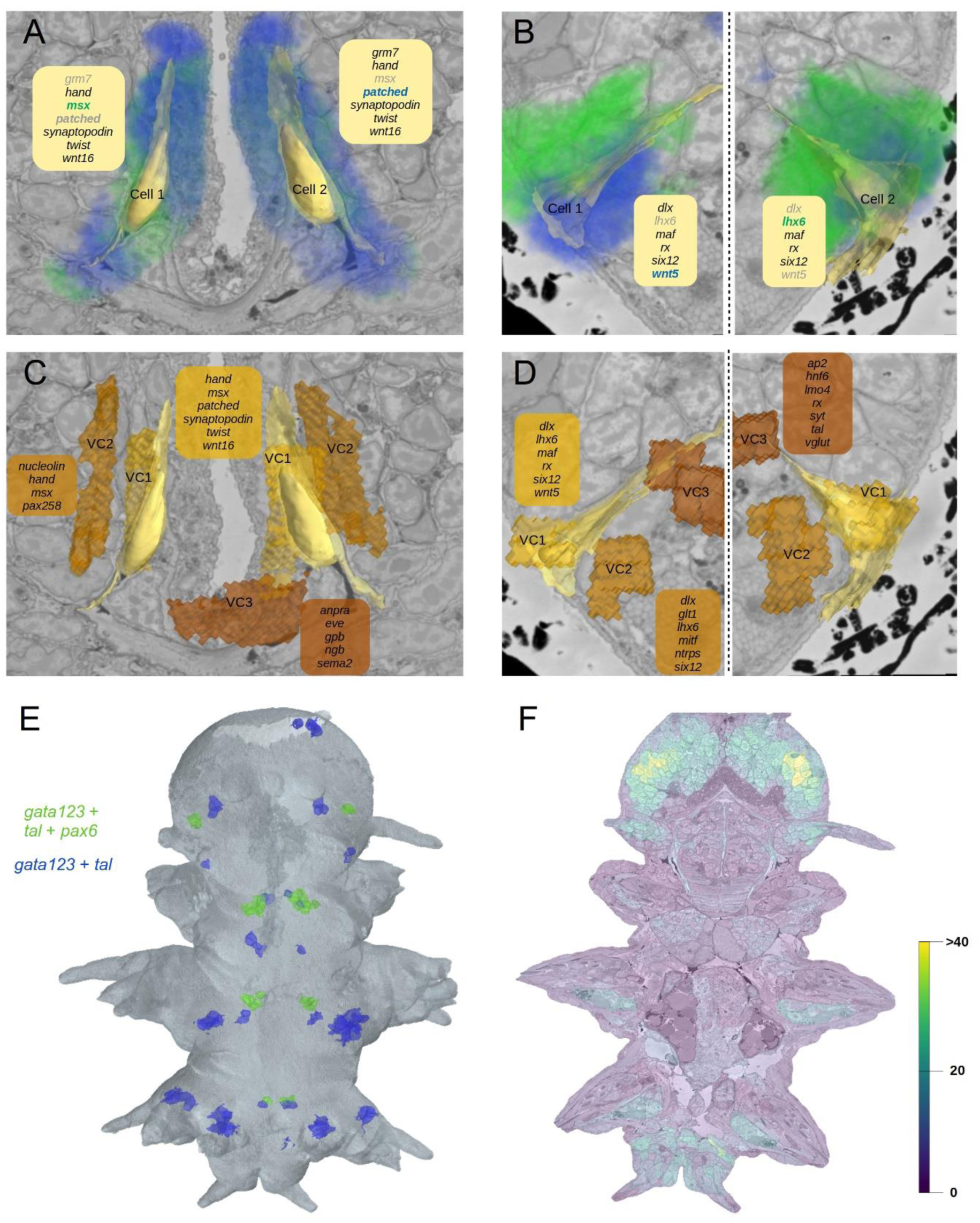
Assigning virtual cells to segmented cells. The difference between assignment by overlap and assignment to genetically nearest Virtual Cell. **A** and **B**: Assignment by overlap: biological variability and registration error resulted in slight asymmetry of the gene expression volumes of the genes *msx* and *patched* (green and blue, A), and *lhx6* and *wnt5* (blue and green, B). Gene lists correspond to genes which would be assigned as expressed in the cell if the assignment was done by volume overlap (regular print for genes with >50% overlap, light grey font for the rest). With a fairly conservative 50% overlap threshold, the resulting assignment for the bilaterally symmetric cells would be different. **C and D**: Assignment to Virtual Cells: for each segmented cell the neighbouring Virtual Cell that has the smallest genetic difference is assigned. This results in a consistent assignment of denoised genetic profiles. **E**: The assignment of the Virtual Cells shown on Figure 3B to the segmented cells. The cells in blue were assigned Virtual Cells that express genes *gata123* and *tal*, the ones in green - genes *gata123, tal* and *pax6*. F: The number of genes expressed in segmented cells after the Virtual Cell assignment. For panels A-D and panel C bookmarks are available in the PlatyBrowser.

To overcome this, we performed the gene expression assignment using Virtual Cells, whose expression profiles were denoised and averaged over multiple regions, and thus are less susceptible to single cell variability and closer to genuine cell type expression profiles. For each segmented cell, we started from the list of genes resulting from the overlap assignment and then chose the Virtual Cell closest in gene expression for assignment (‘virtual cell assignment’; **Figure 6C,D**). Notably, although the spatial location of the assigned virtual cells (**Figure 6E**) is consistent with their spatial location in ProSPr space (**Figure 3B**), only in 39% of cases the segmented cell was actually assigned the spatially closest Virtual Cell. This shows that biological variability combined with the registration error limits the precision of cellular assignment based on expression overlap only.

In order to assess and compare the accuracy of overlap versus virtual cell assignment, we benchmarked both assignment strategies by testing morphologically identifiable symmetric cell pairs for identical gene expression profiles. In total, 208 unambiguous symmetric pairs of segmented cells were marked in the EM volume, covering most tissues (**Supplementary Figure 6D,E**). For each cell pair, gene expression differences were compared after overlap versus virtual cell assignment (**Supplementary Figure 6C**). Virtual cell assignment resulted in a mean discrepancy of 2.0 genes per cell, performing ∼30% better than overlap assignment with 2.8 genes mean discrepancy. Specifically, virtual cell assignment performed better in ∼70% of all symmetric pairs.

Since most of the ProSPr atlas genes were targeting specific tissues, gene coverage is not uniform throughout the animal body (**Figure 3A**). This variability is also reflected in the gene assignment by Virtual Cells (**Figure 6F**). We observed that the assignment quality was higher for cells with better expression coverage (**Supplementary Figure 6B**), and thus expect that the addition of more genes to the atlas will further improve the assignment quality.

### PlatyBrowser: a MultiModal Browser for Platynereis dumerilii

The registered ProSPr and SBEM datasets form a valuable resource containing rich biological information. However, the size of this resource (currently 231 image sources adding up to 2.02 TB lossless compressed image data) poses a challenge to its effective interrogation for scientific discovery. We therefore developed the “MultiModal Browser” (MMB), a Fiji (Schindelin et al., 2012) plugin for interactive browsing of such datasets on a standard computer. We use the MMB to deploy the “PlatyBrowser”, which provides interactive access to our complete *Platynereis* dataset. MMB uses BigDataViewer (BDV) (Pietzsch et al., 2015) for arbitrary plane slicing of volumetric image data. This viewing modality is ideal for the exploration of our SBEM data, which is too dense for volume rendering. In addition, BDV supports simultaneous display of image data of different resolutions, thereby enabling the overlay of ProSPr (0.55 × 0.55 × 0.55 µm^3^) and SBEM (10 × 10 × 25 nm^3^) images (e.g., **Figure 4B-H**). Finally, thanks to lazy-loading from a chunked pyramidal image file format (Pietzsch et al., 2015), even the TB-sized SBEM image can be smoothly browsed on a standard computer connected to a file server.

In order to efficiently explore our resource we designed a user interface for choosing from hundreds of image sources of different modalities as well as for adjusting their display settings (**Figure 7A,D**). The registration of the ProSPr atlas to the SBEM dataset permits to compute a (ProSPr based) gene expression profile at each location in the SBEM space. To enable interactive exploration of this information the MMB provides the possibility to query specific image data at the current location. In the PlatyBrowser, this is used to query all ProSPr images and present the result as both a ranked list and gene expression table (**Figure 7D**).

**Figure 7:**
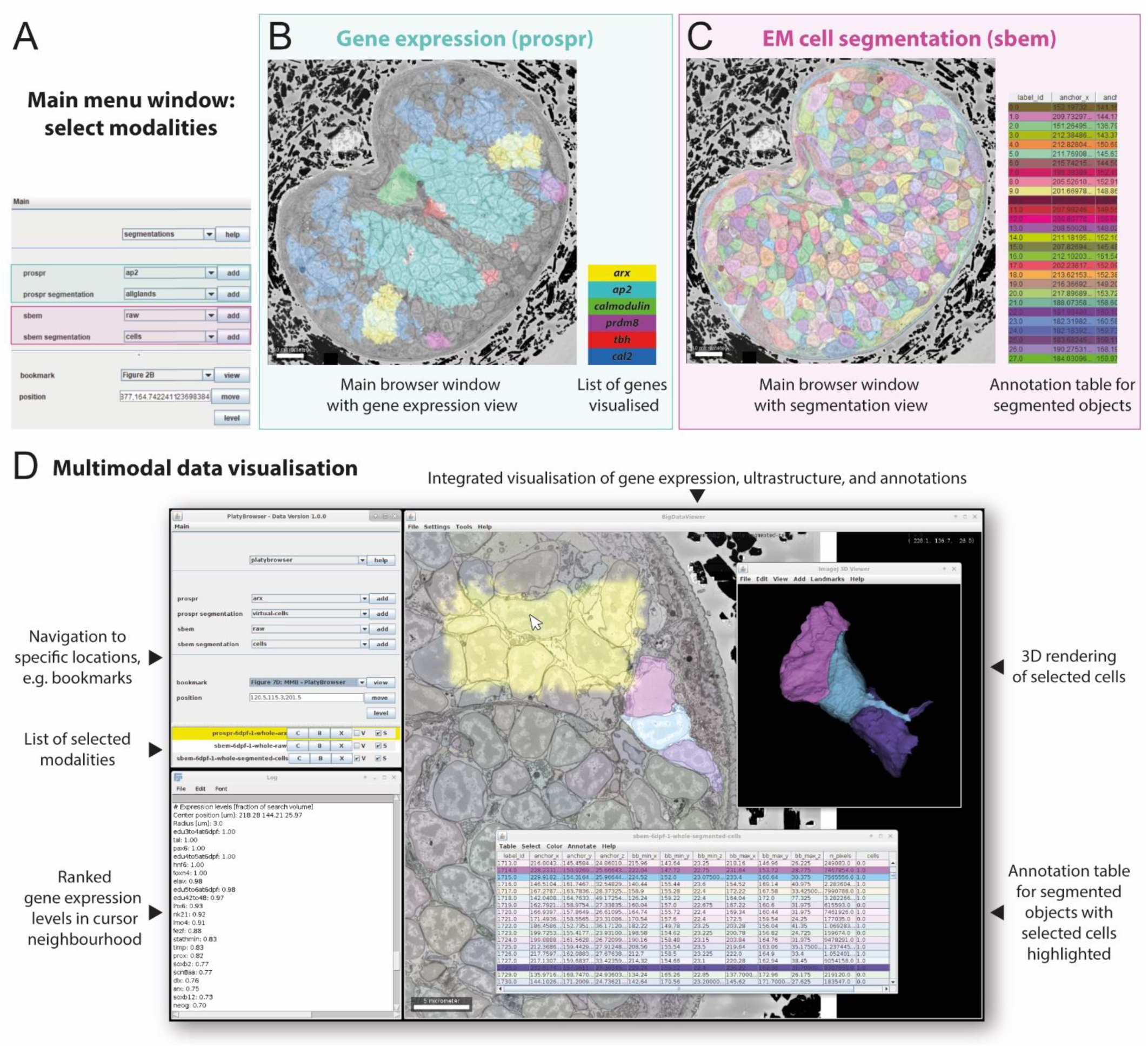
The PlatyBrowser. **A**: User interface to select image sources, change their appearance, and navigate to specific locations in the animal. **B**: BigDataViewer, showing the SBEM image in a region of the adult head, with the ProSPr signal for six different genes. **C**: BigDataViewer of the same section as in B, now displaying the cellular segmentation. **D**: Screenshot of the PlatyBrowser illustrating the integration of modalities and additional functionalities: the expression of gene *arx* is shown in yellow; three segmented neurons are shown next to it; below is shown the annotation table highlighting the rows that correspond to the objects selected. Next to it, the 3D Viewer window shows a rendering of the selected cells; the colours for a given object are identical in the 2D Viewer overlay, 3D rendering and table. Below the main menu, the log window shows a ranked list of the gene expression where the mouse cursor is positioned (white arrow).

To explore segmentations, the MMB provides adjustable lookup tables (Glasbey et al., 2007; Hanslovsky et al., 2020) to overlay each segment with a distinct colour onto the image data. We use this functionality to visualise nuclei, tissue and cell segmentations on top of the image data in the PlatyBrowser (**Figure 7B,D**). This makes it possible to visually observe whether distinct tissue regions correlate with specific ultrastructural morphology. Often, not only ultrastructural morphology but also the 3D shape of anatomical regions is indicative of biological function. We therefore added the possibility to render segmented cells in an interactive 3D viewer (Schmid et al., 2010) (**Figure 7D**).

The nuclei, cell and tissue segmentations allow for the extraction of features such as cellular morphological descriptors or gene expression profiles (**Figure 5** and **Figure 6**). To interactively explore these measurements, the MMB displays a table for each segmentation, where table rows correspond to segmented objects and table columns to feature values. Tables can be sorted according to any feature value and objects of special interest can be selected (highlighted) both in the viewer and in the table. Importantly, the colouring of objects can be configured to reflect any feature value, making it possible to overlay object-based measurements on the image data (see **Figure 5D,E** and **Figure 6F**).

In addition, we implemented a functionality to manually assign object annotations by making the tables editable. For example, we have used this feature to assign the cells in the head to ganglia. We also implemented a bookmark function that not only stores positions and views, but can also update the whole browser state, including segmentations, tables, and objects rendered in the 3D viewer. Bookmarks were used to provide interactive versions for many figure panels presented in this manuscript (see figure legends).

To ensure consistent data access we implemented a versioning system, where the version is selected upon start of the PlatyBrowser. For efficient and open access we host all numerical data on github and stream the image data from a publicly accessible object store using the n5-aws-s3 data format (Pisarev and Saalfeld, 2020). This design allows everyone with an internet connection to browse the TB-sized atlas from a standard computer (for details see Suppl. Material).

## Discussion

### Seeking multimodality

There is a convergence across many fields on the need of integrating distinct modalities to achieve a complete classification of the parts (cell-types) that will hopefully one day guide our understanding of the functioning of animals (Bates et al., 2019; Hobert et al., 2016; Nicovich et al., 2019). In particular, correlating gene expression with cellular and subcellular morphology is crucial to understand the principles that guide the decoding of expression information into cellular phenotypes.

One direction has been to complement expression profiling efforts with cellular imaging. Several recent studies have succeeded to make this link for restricted sets of cells. For example, retrograde labelling of neurons prior to sequencing allows integration of partial projection information with transcriptomics (Kim et al., 2019; Phillips et al., 2019; Tasic et al., 2018). Similarly, photoconversion of dyes in selected cells can be used as a method to correlate single-cell sequencing with live imaging techniques (Lee et al., 2019; Pfeffer and Beltramo, 2017). Other techniques based on probe hybridisation or local sequencing provide expression information *in situ*, recovering the position of the cells interrogated (reviewed in (Lein et al., 2017)). Such spatial transcriptomics techniques can be correlated and complemented with single-cell transcriptomics (Kim et al., 2019; Phillips et al., 2019; Qian et al., 2020), and recent developments using viral barcoding have also enabled the integration of long-range projection information in a brain-wide manner (Chen et al., 2019). However, these approaches only retrieve multimodal information for small subsets of cells and/or their scalability is not straight-forward beyond a set of tissue slices (Ortiz et al., 2019), though this could suffice for small organisms (Ebbing et al., 2018).

The other possible direction is to complement electron microscopy with expression profiling. So far, such attempts have focused on cell type populations (Davis et al., 2020), or have relied entirely on prior knowledge about the cells of interest. For example, this has been accomplished between individuals of the same species for recognisable neurons such as mechanoreceptors, nocireceptors and mushroom body neurons in the *Drosophila* larvae (Ohyama et al., 2015; Saumweber et al., 2018) or peptidergic and ciliomotor neurons and mechanoreceptors in the *Platynereis* larvae (Bezares-Calderón et al., 2018; Verasztó et al., 2017; Williams et al., 2017). This approach can ultimately generate a catalogue of functional, molecular and morphological data for a multitude of cells in an organism, as is the case for the *C.elegans* nervous system with only 302 neurons (Hobert et al., 2016). Recent proposals for the multimodal description of the *Drosophila* central nervous system, with 5.000 times more neurons than *C.elegans*, have been put forward (Bates et al., 2019). The rationale behind these approaches relies on anatomical information gathered through the use of cell-type-specific genetic markers and/or GAL4 lines. This restricts the use of cell type integrative approaches to be done in a cell-by-cell manner, making it hard to scale to the organism level and hard to translate to animal species in which genetic manipulation of cell-types is labour-intensive.

We now push these efforts to a new level, by providing the first multimodal atlas combining electron microscopy and expression data for a fully differentiated specimen. Our success capitalises on two main features of the chosen model species. First, the relatively small size of the 6 dpf *Platynereis* larvae, allows complete acquisition in one SBEM dataset (Titze et al., 2018). Importantly, this stage already comprises hundreds of differentiated cell types and thus displays a maximum of phenotypic information in a small volume. Second, the remarkably high stereotypy of *Platynereis* development, which has enabled the construction of the ProSPr gene expression atlas with cellular resolution for this stage (Vergara et al., 2017), and which we now exploit to register the gene expression to the EM volume.

### A fully segmented volume EM dataset of an entire animal

Ultrastructural analysis of entire animals so far involved tedious collection of manual serial sections (Hall and Altun, 2007; Randel et al., 2014). By adapting SBEM approaches to *Platynereis*, we have achieved the imaging of one full individual at ultrastructural resolution (10 × 10 × 25 nm^3^ voxel size). This volumetric EM dataset enables analyses that span the length scales between the anatomy of entire organs and the ultrastructure of cells and organelles. Adding to this, we have pioneered the detailed comprehensive morphological analysis of such large a EM dataset, which was limited so far by the lack of automated image segmentation methods. Therefore, targeted manual or semi-automated reconstruction by human annotators was used for sparse morphological and phenotypic characterisation (Anderson et al., 2011; Bumbarger et al., 2006, 2013; Cardona et al., 2012; Helmstaedter et al., 2013; Kasthuri et al., 2015; Motta et al., 2019; White et al., 1986). More recently, machine learning-based reconstruction pipelines enabled the segmentation of entire neuronal circuits at synaptic resolution (Berning et al., 2015; Dorkenwald et al., 2017; Heinrich et al., 2018; Januszewski et al., 2018; Li et al., 2019; Staffler et al., 2017; Xu et al., 2020). Extending the pipelines of (Beier et al., 2017; Pape et al., 2019), we have now presented the first automated segmentation of individual cellular somata and nuclei for a complete animal imaged in EM. As a resource, we also provide the code of our algorithms and the weights of the CNNs that we have trained. Our method segments the whole multiple TB dataset in less than three days on a compute cluster and can be directly applied to similar datasets.

### Towards a genetic definition of tissue

A tissue is commonly defined as an ensemble of similar cells and their extracellular matrix with a specific function. Classically, identification of tissues relies on similarity in phenotype of the partaking cells. Taking advantage of our unique dataset with 140 morphological descriptors for more than 10,000 segmented body cells, we provide a first morphometry-based clustering for all cells of an entire body. This leads to the gross subdivision into nervous, muscular, gut, and epithelial cells (**Supplementary Figure 2F)**, which is somewhat expected given the visually discernible size, shape, and/or texture of the cells in these tissues. Unexpectedly, however, we find a brain cluster of neurons that differs in chromatin texture and intensity from all other clusters, hinting at differences in regulatory state. In this context, it is highly remarkable that chromatin features alone are efficient in detecting bilateral cell pairs that are known to share expression profiles (Vergara et al., 2017). This indicates a close link between regulatory state and chromatin morphology.

Our work also highlights the power of using genetic similarity between cells to identify tissues. Previous work identified transcription factors that control tissue integrity for example via the establishment of an ‘adhesive code’, as shown for Pax6, which drives adhesion in the developing retina (Rungger-Brändle et al., 2010). Furthermore, systematic integration of epigenomic and transcriptomic data revealed ‘driver transcription factors’ orchestrating tissue specification, including Pax6 for brain tissue (Zhang et al., 2019). However, these studies were necessarily limited to the genetic characterization of morphologically discernable tissues (requiring dissection, for example for the ENCODE project (ENCODE Project Consortium, 2012)). It has not yet been possible to define the tissues of an animal body purely on genetic grounds. How do genes subdivide the body into coherent tissues?

Towards this aim, recent whole-body single-cell transcriptomics provide genetic clustering of all body cells (Achim et al., 2018; Packer et al., 2019; Sebé-Pedrós et al., 2018a, 2018b). In the absence of spatial mapping at late differentiation stages, however, these studies do not resolve to what extent, and at what hierarchical level, the genetically defined clusters of cells represent coherent tissues. Our genetic clustering of segmented cells based on 201 differentially expressed genes subdivides the *Platynereis* body into 15 clusters. These comprise muscle, midgut, gland, epidermal, and neural cells - representing basic body tissues also recognised by morphological clustering (see above) and by the single-cell transcriptomics studies (Achim et al., 2018; Packer et al., 2019; Sebé-Pedrós et al., 2018a, 2018b). Other clusters are less obvious to interpret without spatial information. First, the neural clusters form two separate larger assemblies, representing ventral nerve cord and brain (**Figure 5B**). Our data do not yet reveal fundamental differences in effector genes between the two, which may change with better gene coverage. Second, the boundaries between the six brain clusters match those between segmented brain ganglia remarkably well. Hence, ganglia represent ensembles of cells that are *genetically* similar, termed ‘transcriptional domains’ (Achim et al., 2018). Our data further show that ganglia are best defined by the combined expression of several regulatory genes rather than by that of a single gene. This includes transcription factors such as Pax6 and other homeodomain factors with high tissue specificity values in our data (**Figure 5D,E**) that have been implicated in tissue specification in other studies (Rungger-Brändle et al., 2010; Zhang et al., 2019).

### MMB and the PlatyBrowser

The MMB makes possible the remote browsing of hundreds of multimodal and multi-scale image data sources. Moreover, not only image data, but also features of segmented objects in tabular form can be visualised and browsed. The MMB is implemented in the popular cross-platform software ImageJ and can be installed by enabling a single Fiji update site (see Suppl. Material), making it an accessible, versatile and extensible platform for browsing image-based atlas data. We expect it to become a valuable tool for exploring such multimodal datasets.

We used the MMB to deploy the PlatyBrowser, a resource for exploration and analysis of the data presented in this publication. The PlatyBrowser currently includes high-resolution SBEM, light microscopy-based gene expression, as well as tissue and cell segmentations. Considering the high degree of stereotypy of *Platynereis*, new image data modalities can be readily added. To this end, we provide all image meta-data and tables within a git-based versioning system to accommodate further improvements of the registration or segmentations, as well as contributions of entirely new imaging modalities from the community. The image data itself is hosted on a web object store for efficient on-demand browsing. The PlatyBrowser is thus an accessible and extendable resource for the investigation of *Platynereis dumerilii* ultra-structural morphology and gene expression.

### Conclusion and outlook

Our work represents the first step towards a richer interrogation of the relationships between transcriptomics and subcellular morphology. For example, our resource will further improve with the addition of new genes into ProSPr. Moreover, single cell transcriptomics data can be spatially mapped into the ProSPr atlas (Achim et al., 2018) and thus onto the segmented EM cells. For this, we can capitalise on the assignment of Virtual Cells to segmented cells as an intermediate for combining expression space and morphospace, given that it generates denoised gene profiles devoid of gene expression leaking from neighbouring cells. Next, the segmentation of additional ultrastructural features (e.g. mitochondria, Golgi apparatus) in our dataset will allow novel interrogation of correlations between genotype and phenotype. We also envisage that enhanced resolution of new EM datasets registered onto our resource will allow the automated reconstruction of neuronal circuits and mapping of synapses at larger scale. Notably, a complete serial section EM dataset already exists for an earlier stage (3 dpf) of *Platynereis*, and the neuronal sensory-motor circuits mediating visual navigation (Randel et al., 2014; Verasztó et al., 2018), ciliary swimming (Verasztó et al., 2017) and startle response (Bezares-Calderón et al., 2018) have already been traced. This provides a catalogue of cell types based on dendritic and axonal morphology (Williams and Jékely, 2019) some of which can be linked to expression data, as illustrated for tens of cells in the neurosecretory center of Platynereis at 3 dpf (Williams et al., 2017). With our new resource, this can now be extended to include connectomics and transcriptomics information for an entire animal, and for different developmental stages. Additionally, using the multimodal capacity of the PlatyBrowser, we will be able to incorporate physiological data for identified cell types using tools such as calcium imaging (Chartier et al., 2018) and Crispr-cas9 (Bezares-Calderón et al., 2018). With the resources that we have developed for *Platynereis*, we expect this approach to be suited for other animals. The stereotypy required for the registration of multiple individuals is not unique to *Platynereis* and can be found within many metazoan lineages. This is the case for model organisms such as nematodes (Schafer, 2016), *Drosophila* (Jenett et al., 2012), *Aplysia* (Katz and Quinlan, 2019), the chordate *Ciona* (Satoh, 1999), and early neurons of zebrafish (Metcalfe and Westerfield, 1990). Importantly, the experimental techniques used for data collection (electron microscopy, *in situ* hybridization, single cell RNA sequencing) are not species-specific and can be readily applied to non-canonical laboratory species. The complete integration of whole-body connectomics and transcriptomics can bring back more diversified studies of the animal kingdom, using clearing methods if needed to build atlases based on light microscopy. This will open the door to the *in toto* comparison of cell types and neural circuits within and across organisms at the genetic and ultrastructure level, bringing us closer towards the understanding of the function, development and evolution of nervous and other systems.

## ACKNOWLEDGEMENTS

We want to thank Gemma Estrada Girona for assembling Figure 7 and for input in Figure 4. Thanks to the IT department at EMBL for setting up the cloud infrastructure, esp. Josep Manel Andres Moscardo. Many thanks to Stephan Saalfeld, Igor Pisarev and esp. Tobias Pietzsch for their help with software design. Thanks to Giulia Mizzon for providing annotations for the segmentation validation, and David Puga for annotating points for construction of the *Platynereis* midline. HMV was supported by an EMBO Long Term Fellowship.

## AUTHOR CONTRIBUTIONS

<u>Conceptualization</u>: AK, CT, DA, HMV, YS; <u>Methodology</u>: AAW, AK, CG, CP, CT, HMV, KM, VZ; <u>Software</u>: AAW, BT, CP, CT, HMV, KM, VZ; <u>Validation</u>: CP, CT, DA, HMV, KM, PM, PYB, RT, VZ, YS; <u>Formal analysis</u>: CP, HMV, KM, OS, VZ; <u>Investigation</u>: BT, CG, ELS, HMV, OS, PM, PYB; <u>Resources</u>: AK, CP, CT, HMV, KM, VZ; <u>Data curation</u>: CP, KM, VZ; <u>Writing - original draft</u>: AK, CP, CT, DA, HMV, KM, RT, VZ, YS; <u>Writing - review and editing</u>: AAW, AK, BT, CG, CP, CT, DA, HMV, KM, RT, RWF, VZ, YS; <u>Visualization</u>: CP, CT, HMV, KM, RT, VZ, YS; <u>Supervision</u>: AK, CP, CT, DA, HMV, YS; <u>Project administration</u>: DA, YS; <u>Funding acquisition</u>: AK, DA, RWF, YS

## DISCLOSURE DECLARATION

AAW is founder and owner of ariadne-service gmbh

## Materials and Methods

### Sample fixation, preparation and imaging

6 dpf *Platynereis dumerilii* were anaesthetised using 7% MgCl_2_ in seawater (1:1 ratio). Platynereis was fixed in a solution of 2% formaldehyde and 2.5% glutaraldehyde in 0.1M sodium cacodylate buffer for 4 days at 4 degrees Celsius.

Fixed samples were prepared for SBEM following an adapted form of the NCMIR protocol (Deerinck et al., 2010) aided by microwave application (Pelco Biowave). Samples were postfixed with 2% osmium tetroxide in 1.5% potassium ferrocyanide (14 mins of 2 min on/off cycles, 150W, with vacuum) followed by rinsing with H_2_O. The rinse protocol, used throughout processing, involved one initial exchange of H_2_O on the bench and twice aided by the microwave (40 seconds, 80W). The samples were then incubated in 1% aqueous solution of thiocarbohydrazide (14mins of 2min on/off cycles, 150W, with vacuum with the cold spot set to 40 degree celcius), followed by rinsing with H_2_O. This was followed by a second step of osmium tetroxide, this time in a 2% aqueous solution (14 mins of 2 min on/off cycles, 150W, with vacuum) and another rinse step. Samples were then incubated in 1% uranyl acetate (aqueous) overnight at 4°C. The following day the samples were rinsed and the final step of staining was performed. Samples were transferred to Walton’s lead aspartate solution for 14mins in the microwave (2min on/off cycles, 150W, vacuum, 50°C). Samples were again rinsed with H_2_O and then dehydrated with increasing concentrations of ethanol (20%, 50%, 70%, 90%, 3 × 100%). Samples were then infiltrated with durcupan resin through increasing percentages of resin with ethanol (25%, 50%, 75%, 3 × 100%). Samples were first prepared using the minimal resin method (Schieber et al., 2017) and then embedded in silver epoxy resin (Wanner et al., 2016).

Samples were mounted onto aluminium pins with 2-part silver epoxy. Images were acquired with a ZEISS Merlin SEM at 1.8 keV landing energy, 270 pA beam current, and 0.8 µs dwell time. ∼2.5 TB comprising 11,416 slices with >200,000 image tiles were acquired at 10 × 10 nm^2^ pixel size and 25 nm cutting thickness. For acquisition we used the open-source acquisition software SBEMimage (Titze et al., 2018). For image registration, translational offsets between neighbouring image tiles were calculated using a custom optimised normalised cross-correlation procedure. Subsequently, offsets were used to optimise the tile positions in a global total least square displacement sense. Before in-plane stitching, the histograms of neighbouring image tiles were matched in order to adjust and homogenise the contrast and brightness. A subset of about 10% of the sections showed nonlinear distortion artefacts due to sample charging. For these sections, a non-distorted neighbouring section was manually chosen as a reference for distortion correction using the ImageJ plugin bunwarpJ (Arganda-Carreras et al., 2006).

Intensities of the fully stitched z-slices were then matched to remove intensity and contrast jumps in the z-direction. This was done by calculating the 5% (L) and 95% (U) quantiles of intensity for each slice within the *Platynereis* (excluding the resin and silver embedding) from a downsampled version of the raw data (pixel size of 0.32 × 0.32 × 0.025 microns). The 95% quantile U was matched between slices to adjust for shifts in absolute intensity, while the quantile range U - L was matched between slices to adjust for shifts in contrast. For each slice, these adjustments can be written as a linear transformation of the intensities: x_c_ = ((U-L)_ref_/(U-L)) (x–U) + U_ref_ where x_c_ are the corrected intensities of the slice, x are the raw intensities of the slice and ‘ref’ refers to the reference which was taken as the median of all slices. Using the median intensity of the resin around the *Platynereis* as a reference (as this should be fairly constant throughout the dataset), we noticed that this technique performed less well at the start and end of the dataset, where only the tips of the *Platynereis* remain. To correct for this, we chose a minimum z cutoff of slice 800 and a maximum z cutoff of slice 9800 – values beyond these cutoffs were corrected by the median of the correction factors of the 100 slices next to each cutoff (i.e. the calculated values are extrapolated for the very tips of the dataset).

Scripts for the intensity correction in z can be found on github: https://github.com/platybrowser/platybrowser-backend/tree/master/misc/intensity_correction

### Segmentation Methods

In total, we provide segmentations of all cells, all nuclei, the cuticle and selected tissues and body parts as well as nuclear chromatin. For simpler segmentation tasks (tissues and regions of the animal: coelomic cavity, glands, gut, secretory cells and yolk) where the region boundary is very pronounced we use the carving workflow of ilastik (Berg et al., 2019) on downsampled data (80 × 80 × 100 nm). Similarly, to segment nuclear chromatin we use ilastik pixel classification, limiting it to the pre-segmented nuclei regions. For ilastik training the nuclei of 50 cells with diverse nuclear morphology were interactively annotated as “heterochromatin and nucleolus” or “euchromatin” classes, on data downsampled to 20 × 20 × 25 nm voxel size.

For the more complex tasks of cell and nuclei segmentation as well as for the segmentation of the cilia and cuticle, we extended the state-of-the-art EM segmentation methods originally developed for neural tissue blocks. In essence, the pipeline consists of a membrane detection step performed by a 3D U-net and a graph agglomeration step performed either by the Lifted Multicut or the Mutex Watershed algorithms.

In detail, we start from the segmentation of the nuclei. The groundtruth annotations for CNN training were provided by ariadne.ai (12 blocks of 400 × 400 × 120 pixels each) and additionally curated. A 3D U-net was trained to predict short- and long-range pixel affinities as described in (Lee et al., 2017) and to predict for each pixel whether it belongs to a nucleus. The predictions were processed by the Mutex Watershed algorithm (Wolf et al., 2018), blockwise in blocks of 512 × 512 × 64 pixels. We chose Mutex Watershed over the more common superpixel-based Multicut agglomeration as we observed Multicut frequently merges individual nuclei which touch across a very small portion of their boundary (short-circuiting of the multicut constraints). The individual block segmentations were then stitched together using Multicut-based agglomeration with edge weights derived from pixel affinites as described in (Pape et al., 2017). All computations were done on raw data downscaled to the resolution of 80 × 80 × 100 nm.

Cilia and cuticle segmentation were performed using the same method as for the nuclei. The cilia segmentation was performed at full resolution (10 × 10 × 25 nm^3^ voxel size) but only applied to the segmented nephridia cells, using 3 blocks of annotated training data consisting of a total of 171 megavoxel. The cuticle segmentation was performed for data downscaled to 40 × 40 × 50 nm using 5 training blocks consisting of a total of 495 megavoxel.

Cell segmentation was also started from membrane detection. The groundtruth annotations were provided by ariadne.ai, consisting of 8 blocks of 628 × 628 × 130 pixels that were additionally curated and extended by one additional block of size 1280 × 1280 × 120 pixels to include more biological variability. A 3D U-net was trained to predict short- and long-range pixel affinities. In addition, we insert the edges of tissue and region segmentations (see above) into the affinity predictions, in order to avoid missing boundary signal due to the very different appearance of some region/tissue boundaries. These predictions were then used to break the volume into superpixels by the blockwise distance transform-based watershed algorithm (Beier et al., 2017). The superpixels were used to construct a region adjacency graph and to solve the segmentation problem as a graph partitioning with Lifted Multicut (Horňáková et al., 2017). Unlike (Beier et al., 2017) and other connectomics pipelines, we additionally exploit the nuclei segmentation to enforce separation of cells containing different nuclei. To that end we introduce lifted edges between superpixels which belong to the segmented nuclei, attractive for the superpixels of the same nucleus and repulsive for the superpixels of different nuclei. Lifted edges are introduced up to a graph distance of 4 and the attractive / repulsive edge weight is set to the maximum / minimum of the local edge weights. The overall lifted multicut problem was solved by the hierarchical solver introduced in (Pape et al., 2019). Since the nuclei repulsion is only included up to a certain graph distance, there are still objects in the resulting cell segmentation that contain more than one nucleus segment. We find these in post-processing and separate them individually by running a graph watershed seeded from the nodes mapped to the nuclei. Cell segmentation was performed on the raw data downscaled to a voxel size of 20 × 20 × 25 nm^3^; the runtime for the whole volume measured 10 hours on 6 GPUs for the neural network prediction and 20 hours on a CPU compute cluster for the agglomeration part.

Besides tissues listed above, several ganglia in the animal’s head were composed by manually selecting the corresponding segmented cells. We also subdivided the body into its parts

Our motivation to include prior information on the nuclei comes from the observation that the 3D U-net does not generalise sufficiently well to the diversity of cell boundary appearance in the whole organism volume. While the lack of generalisation can be addressed by creating additional training data, for a 3D EM segmentation task this process is extremely laborious. Instead, we prefered to rely on the nuclei segmentation - a much simpler problem which our algorithm solves to 99.0% accuracy and enforce the constraint for every cell to only contain one nucleus which at this developmental stage is expected to be true for the absolute majority of cells.

The final proof-reading was performed in a semi-automated manner. First, we compute a morphology-based score for all cells, that roughly matches the likelihood of a cell being a false merge. Then, we iterate through the top 1000 cells based on their rank by this score and correct all cells that contain a false merge by running graph watershed from user-generated seeds. This resolving step had to be applied to 154 falsely merged cells. In addition, we use Paintera (Hanslovsky et al., 2020) to perform some more fine grained proof-reading. Note that the segmentation currently provided in the PlatyBrowser has not been proof-red down to the pixel level of every cell. We corrected the errors we found by the approach above and additionally polished the segmentation of the regions used in the analysis presented here: nephridia, adult eyes, symmetric cells used for the gene assignment validation. For reference, finalising the segmentation of the nephridia (14 cells with far-reaching cilia) from the fully automatic pipeline results took approximately 2 hours with Paintera.

In addition, semantic segmentations based on a CNN for the muscles and the neuropil were provided by ariadne.ai.

We provide the weights for the networks used to segment cell membranes, cilia, cuticle and nuclei on zenodo https://doi.org/10.5281/zenodo.3675288 as well as the corresponding training data https://doi.org/10.5281/zenodo.3675220. The ilastik project and training data for the chromatin segmentation is also available at https://doi.org/10.5281/zenodo.3676534. The ilastik projects for carving the animal outline and regions/tissue are available at https://doi.org/10.5281/zenodo.3678793. The scripts to run the segmentation methods are available at https://github.com/platybrowser/platybrowser-backend/tree/master/segmentation.

#### Segmentation Validation

The validation of the segmentation was based on manual annotations for cell centers and nuclei on 8 slices (4 transversal, 4 horizontal) from 8 domain experts, each slice annotated at least twice without access to the automatic segmentation results. The missing detections in the expert annotations were then additionally corrected by comparison with the automatic segmentation.

In total we observe, for the cell segmentation task, 6.45 % false split errors and 3.26 % false merge errors, based on 4806 annotations. For the nucleus segmentation task, we found 0.49 % false positive detections; 0.55 % false negative detections based on 2888 annotations.

**Supplemental Figure 2A** gives an overview of the expert annotations and 2 examples of false merge and false split errors each.

### Foregut cDNA library

Around 40 maturing animals (juveniles) were isolated from the tubes and starved for three days. The epidermis around the foregut was removed to reveal the muscular pharynx and the anterior part of the organ. The tissue was immediately fixed in liquid nitrogen. Tissue was then ground in Trizol and a Trizol/Chloroform extraction was performed. RNA was purified with RNeasy kit (Qiagen columns). Illumina library was prepared by the EMBL GeneCore facility. cDNA for individual gene cloning was generated with SuperScript III kit (Invitrogen).

### In situ hybridisation and generation of gene expression maps

Animal breeding, animal fixations, and whole-mount *in situ* hybridisations were performed as described previously (Tessmar-Raible et al., 2005). Light microscopy sample imaging and gene expression maps were generated following the ProSPr protocol as described in (Vergara et al., 2017).

### Generation of virtual cells

The generation of Virtual Cells (VCs) was done as in (Vergara et al., 2017) with some modifications (see also main text). Analysis of ProSPr data and generation of VCs was done using the software R Bioconductor (Gentleman et al., 2004). The atlas was binned by a factor of 3 to generate supervoxels (SVs). Each SV represents a cube of 27 pixels (4.5µm^3^ total volume). On average, and given the size of a nucleus in the EM dataset (67.32 +/- 15.19 µm^3^), each cell is minimally covered at least by 12 SVs. Note that we used the nucleus size to estimate cell coverage as the big majority of cells are neurons (small cytoplasm), and supervoxels along cell boundaries will result in noisy signals. Next, we generated a matrix with the SV spatial location in xyz and the proportion of pixels within that SV positive for each gene. SVs with very low correlation with their neighbours were removed from the analysis as they represent noise. Due to memory constraints, the SVs were subdivided into different anatomical regions (e.g. head, ventral nerve cord, foregut, etc) for further analysis.

Next, SVs were grouped into Virtual Cells (VCs). This is done by a recursive process of hierarchical clustering to find groups of SVs based on their similarity in expression profile. This process does not take into account the location of the SVs, but it is constrained by the final size of the group, allowing groups to be formed so that they represent between 1 and 6 cells, always optimising for expression correlation. For hierarchical clustering we used the function ‘hclust’ with the method *‘complete’*. For distance calculation we used the method *‘jaccard’* from the *‘vegan’* package. For computing SVs correlation we used the basic R function ‘*cor*’ with the following parameters: *‘use = “complete.obs”, method = “kendall”’*.

In contrast to (Vergara et al., 2017), VCs were automatically curated based on size and spatial distribution of their constituent SVs. In more detail, first all the VCs that contain less than 16 supervoxels were filtered out, since below this size they consistently showed low spatial correlation (probably representing cell boundaries). Next, VCs were split into spatially connected components (CCs), and the ones below size 5 supervoxels in size were filtered out (cell size values below 5 SVs do not properly represent a cell). All CCs comprised of 8 or more supervoxels were kept without further checks, and the ones in the range of [5,8) were checked for a symmetric partner - if there was another CC of the same VC symmetrically located on the other side of the animal, the CC was considered plausible and retained. The partner CC was considered symmetric if located within the radius of 4 supervoxels (one cell diameter) from the mirrored coordinates of the centre of the given CC. After this curation step the VCs are assembled together again (so they are retained as bilateral cell types) and their expression profile is calculated.

The complete automated pipeline for generating Virtual Cells from the ProSPr gene expression maps can be found on GitHub: https://github.com/platybrowser/prospr-vc-generation.

### Registration of ProSPr to EM

We used the software package elastix (Klein et al., 2010; Shamonin et al., 2013) to compute a multi-step registration of an average DAPI signal (representing nuclei of 153 specimens from the ProSPr resource) onto the binary mask of the segmented nuclei of the EM individual. Historically, elastix is mainly used by the medical image analysis community. To increase the usability of elastix for the biological community we developed the Fiji (Schindelin et al., 2012) plugin ElastixWrapper (Tischer, 2019), which we used for the registration. As an input for the registration we prepared three image files at an isotropic voxel spacing of 0.55 µm: DAPI: a greyscale image of the average DAPI signal of ProSPr (Vergara et al., 2017); EM-Nuclei: a binary image containing the segmented EM nuclei; and EM-Mask: a binary mask covering the image region containing information relevant for the registration optimisation (**Supplementary Figure 4**).

To prepare the EM-Nuclei image we removed 42 nuclei in the gut region from the nuclei segmentation binary mask as these nuclei do not locate to stereotypical positions and were not reflected in the ProSPr DAPI signal, and therefore they did not properly guide the registration. Next, we downsampled the binary mask of the remaining 11,456 segmented nuclei to the 0.55µm voxel size of the ProSPr data; note that due to the bi-linear downsampling algorithm the voxel values in the EM-Nuclei image are not strictly binary, but show some shades of grey (**Supplementary Figure 4B**).

To prepare the EM-Mask image we isotropically dilated the EM-Nuclei image by 12 pixels, corresponding to 6.6 µm. The EM-Mask serves to restrict the optimisation algorithm of elastix to relevant regions of the image. The usage of such a mask in elastix is optional, we however found it critical for our image data, because it contains a lot of empty space due to the oblique orientation of the EM image within the 3D voxel space.

Using these images we sequentially ran elastix with transformation models of increasing deformability: Similarity: a rigid transformation, allowing for rotation, translation, and a uniform scaling factor; BSpline100: a locally deformable BSpline transformation (Rueckert et al., 1999) with a 3-D voxel grid spacing of 100 pixels (55 µm); BSpline30: BSpline transformation with a grid spacing of 30 pixels (16.5 µm); BSpline10: BSpline transformation with a grid spacing of 10 pixels (5.5 µm). The registrations build on each other, e.g. the BSpline100 transformation takes the Similarity transformation as a starting point. Importantly, since the two data sets are initially very misaligned (**Supplementary Figure 4A,B**) using the automated registration algorithms of elastix directly did not work for us. The reason is that elastix uses local search algorithms for improving the transformation parameters and thus typically needs a decent starting condition. To produce such a starting condition, we used the Fiji plugin TransformJ Rotate (https://imagej.net/ImageScience) to manually determine angles that roughly align the two data sets in 3-D. We then converted these angles into an elastix transformation file (Manual-Rotation). Using the Manual-Rotation transformation as a starting point, we could then use elastix to sequentially improve the registration. After each step in the registration sequence we visually compared the transformed DAPI image with the EM-Nuclei image to qualitatively assess whether the registration improved (e.g., using overlays as shown in **Figure 4A** in the main text).

To further assess registration quality, we manually identified 43 corresponding landmarks between the EM and ProSPr datasets, covering all relevant regions of the specimen (**Supplementary Figure 3D**). These landmarks were selected based on unequivocal positions in both datasets using the nuclei, muscle and neuropil signals. Using these landmark pairs we could measure the registration quality at each step by transforming the manually assigned landmark coordinates in EM space into ProSPr space and then comparing these computed coordinates with the manually assigned coordinates. Measuring the pairwise distances of the 43 coordinate pairs for the different registration steps we obtained the following results (mean, median): Manual-Rotation: 63.57µm, 58.79µm; Similarity: 11.90µm, 10.09µm; BSpline100: 5.29µm, 4.48µm; BSpline30: 4.23µm, 3.53µm; BSpline10: 3.47µm, 2.99µm. The fact that the mean and median are similar shows that there are no severe outliers and thus the registration quality is consistent across all parts of the animal (**Supplementary Figure 3D,E**). In addition, the pairwise distances decrease with each registration, showing that the elastix algorithm indeed converged to a better registration. We did not attempt to improve the registration beyond the current measured accuracy of ∼3µm, because at 6 dpf the typical variation in the position of individual cells in *platynereis* is ∼4.7µm (Vergara et al., 2017).

Scripts and transformation files used for the registration can be found on github: https://github.com/platybrowser/platybrowser-backend/blob/master/registration/0.6.3

### Morphology clustering

Various morphological, intensity and texture features were calculated from the cell, nucleus and chromatin segmentations (full description of features in **Supplementary Table 2**). Cells (and their associated nuclei and chromatin) were filtered to remove those most likely to be affected by segmentation errors – cells were removed that had no assigned nucleus, were in certain regions (yolk, neuropil, cuticle, cavities), or outside a reasonable size range. This resulted in 11368 remaining cells. All features were calculated on data downsampled to 80 × 80 × 100 nm resolution.

For downstream analysis, any cells (and associated nuclei and chromatin) that were within the region for extrapolated intensity correction (see Sample fixation, preparation and imaging methods section) were also discarded (leaving a total of 10346 cells). This was to avoid any possible artefacts in features that rely on the raw intensity data.

This resulted in a table of 140 features by 10346 cells. All features were standardised by centring to a mean of 0 and scaling to unit variance. A K-nearest neighbour graph was then constructed (k = 10) using Euclidean distance between the feature vectors. Community detection (using the Louvain method (Blondel et al., 2008)) was performed with a resolution parameter of 1.2, resulting in 12 clusters. Clusters were visualised on a UMAP (McInnes et al., 2018) (Uniform Manifold Approximation and Projection for Dimension Reduction) using the following parameters: n_neighbours=10 and min_dist=0.1.

All the morphological clustering analysis was performed in Python with scikit-image (van der Walt et al., 2014), vigra (http://ukoethe.github.io/vigra/), scipy (Virtanen et al., 2020), mahotas (Coelho, 2012), scikit-learn (Pedregosa et al., 2012), networkx (Hagberg et al., 2008), python-louvain (https://github.com/taynaud/python-louvain), umap-learn (McInnes et al., 2018), pandas (McKinney, 2010), numpy (van der Walt et al., 2011) and snakemake (Köster and Rahmann, 2012). All code is freely available in the github repository (the clustering code as a snakemake workflow). The snakemake workflow also contains code for various other analyses e.g. clustering of different subsets of morphological features, plotting of *Platynereis* regions on the UMAP, plotting gene expression on the UMAP, or construction of various heatmaps.

The script for calculation of morphological features is available here: https://github.com/platybrowser/platybrowser-backend/blob/master/mmpb/extension/attributes/morphology_impl.py and the Snakemake workflow for clustering analysis is available here: https://github.com/platybrowser/platybrowser-backend/tree/master/analysis/morphology_clustering

### Bilateral pair analysis

To assess how different subsets of morphological features (**Supplementary Table 2**) perform on finding bilateral pairs of cells in the dataset, we established a set of xyz criteria for being bilateral based on the position of cell’s nuclei. As the *Platynereis* is bent in the EM image volume, we cannot directly use the xyz position of the nuclei to determine if they are bilateral or not. Instead, we first calculated a midline surface for the *Platynereis* by fitting a second order polynomial of two variables to a set of manually chosen points that lie on the midline. The absolute distance from this surface could then be calculated for all nuclei. While this accounts for the distance from the midline, it doesn’t assess if nuclei have a similar position along the anterior-posterior (AP) or dorsal-ventral (DV) axis of the *Platynereis*. To account for this, we transformed the xyz position of all nuclei from the EM space back to the original ProSPr space via the program elastix (Klein et al., 2010; Shamonin et al., 2013) (same transformation parameters as calculated for the registration – see Registration of ProSPr to EM methods section). As the AP and DV axes are nicely aligned to y and z in the ProSPr space, we can use these coordinates as estimates for AP and DV position. The full criteria are then: similar absolute distance from midline; on opposite sides of the midline; similar y position in ProSPr space; similar z position in ProSPr space.

To assess a reasonable range to consider ‘similar’, these statistics were calculated for a set of manually curated bilateral pairs of cells (203 pairs total). The acceptable difference was then set to the mean over these pairs plus two standard deviations.

Given these criteria to assess if cells are bilateral, we then wished to calculate the distance in morphology space that must be travelled from one cell to find its potential bilateral partner. This was calculated first by taking a certain subset of the morphology criteria (calculated as in the Morphology clustering methods section) and standardising them – by centring each feature’s mean to 0 and scaling to unit variance. For each cell, a ranking was then formed (based on Euclidean distance in morphology space) of every other cell in the dataset - from its very closest neighbour, to its most distant neighbour. Then, using the xyz criteria above, the closest neighbour was found in this ranking that was a potential bilateral partner.

To compare to random assignment of bilateral pairs, the ranking of cells in morphology space was randomly shuffled, and the same analysis performed. This was repeated 100 times, and the mean of all trials taken for the final randomised results shown in **Supplementary Figure 2B**.

Scripts for this analysis are available here: https://github.com/platybrowser/platybrowser-backend/tree/master/analysis/bilateral_pairs, and the midline calculation is available here: https://github.com/platybrowser/platybrowser-backend/tree/master/analysis/midline. The midline fit and final graphs were calculated in R, making use of the tidyverse (Wickham et al., 2019) and rgl (https://cran.r-project.org/web/packages/rgl/index.html) packages. All other analysis was in Python with pandas (McKinney, 2010), numpy (van der Walt et al., 2011), scikit-learn (Pedregosa et al., 2012) and scipy (Virtanen et al., 2020).

### Gene expression clustering

Overlap assignment was used to assign a vector of gene overlap values to each segmented cell. In brief, the fraction of the volume of each segmented cell overlapping with the registered volume of every gene was calculated. This resulted in a vector of length 201 (equal to the number of genes) with values ranging from 0 (no overlap) to 1 (complete overlap).

Segmented cells were then filtered to remove those most likely to be affected by segmentation errors – cells were removed that had no assigned nucleus, were in certain regions (yolk, neuropil, cuticle, cavities), or outside a reasonable size range. In addition, any cells that expressed no genes were discarded. This resulted in 11,366 remaining cells.

A K-nearest neighbour graph was then constructed (k = 20) using Euclidean distance between the gene overlap vectors. Community detection (using the Louvain method (Blondel et al., 2008)) was performed with a resolution parameter of 1.2, resulting in 15 clusters.

Clusters were visualised on a UMAP (McInnes et al., 2018) (Uniform Manifold Approximation and Projection for Dimension Reduction) using the following parameters: n_neighbours=20 and min_dist=0.1. All the gene expression clustering analysis was performed in Python with scikit-learn (Pedregosa et al., 2012), networkx (Hagberg et al., 2008), python-louvain (https://github.com/taynaud/python-louvain), umap-learn (McInnes et al., 2018), pandas (McKinney, 2010), numpy (van der Walt et al., 2011) and snakemake (Köster and Rahmann, 2012). The code is freely available as a snakemake workflow: https://github.com/platybrowser/platybrowser-backend/tree/master/analysis/gene_clustering. The snakemake workflow also contains code for various other analyses e.g. clustering of different kinds of gene expression (binarised by a threshold, assignment by virtual cells etc), plotting of *Platynereis* regions on the UMAP, plotting morphology statistics on the UMAP, or construction of various heatmaps.

### Ganglia specificity analysis

Segmented head ganglia were displayed on the same UMAP as the gene expression data (see gene expression clustering methods).

A specificity score was calculated for every gene cluster and individual gene for every ganglia. This specificity score is a combination of two measures: firstly, the fraction of expression confined to a given ganglion (A) and secondly, the fraction of that ganglion covered (B). In analogy to the F1 score, specificity is calculated as: 2AB / (A + B)

The calculation for individual genes relies on overlap assignment to segmented cells (see gene expression clustering methods). A threshold of 0.5 was used to label a cell as expressing a particular gene.

The specificity calculations are part of the same workflow as described in the Gene Expression Clustering section: https://github.com/platybrowser/platybrowser-backend/tree/master/analysis/gene_clustering, specific script here: https://github.com/platybrowser/platybrowser-backend/blob/master/analysis/gene_clustering/scripts/ganglia_specificity.py

### Virtual cell assignment

In order to assign the gene expression patterns to the segmented cells the Virtual Cells (VCs) volume was registered onto the EM one. To compensate for biological variability and registration error for each cell we considered all the VCs found in the radius of 5 µm from the cell boundaries. The expression pattern of the cell calculated by gene overlap was compared to the expression pattern of every VC found. Finally, the segmented cell was assigned to the genetically closest VC. Considering the fact that the available 201 genes were mostly targeting specific tissues, the animal can and does have areas without or with minimal gene coverage. That is why in case for a given cell either no VCs were found around, or an absence of any expression was genetically closer than any of the VC’s found, the cell was allowed not to be assigned any gene expression or, more precisely, to be assigned zero expression for all the genes.

To compare this type of assignment to purely spatial assignment we also assigned the Virtual Cells by overlap: each segmented cell was assigned either the VC that overlapped its volume the most, or no expression in case no VC overlap was present. Such assignment gave a worse result in the symmetric cells test: the mean gene distance between symmetric cells was 3.62 (compared to 1.96 distance for assignment by genetic distance described above). The agreement of these two methods was checked on the segmented cells that contained nuclei only.

### Data provision

#### EMPIAR and BioStudies

Gene expression maps, prospr segmentations and registration files are archived and can be downloaded from BioStudies (https://www.ebi.ac.uk/biostudies/) under the accession id S-BIAD14. Electron microscopy data and EM based segmentations are archived and can be downloaded from EMPIAR (https://www.ebi.ac.uk/pdbe/emdb/empiar/) under the accession id 10365.

#### S3 object store

In addition to EMPIAR and BioStudies we host all image data on an aws-s3 (https://aws.amazon.com/s3) compatible object store at EMBL Heidelberg using the open source implementation of MinIO (https://min.io). The data is stored in the n5 data format (https://github.com/saalfeldlab/n5) which allows chunking and compression for efficient data access. It can be accessed on demand through the n5-s3 API (https://github.com/saalfeldlab/n5-aws-s3).

#### Github

The tabular data derived from images and segmentations is hosted on github: https://github.com/platybrowser/platybrowser-backend/tree/master/data/1.0.0/tables. The repository https://github.com/platybrowser/platybrowser-backend also contains the image metadata in the BigDataViewer xml file format, which serves as an entry point to start the PlatyBrowser for the data hosted on the s3 object store. In order to keep track of changes in the derived data, for example due to segmentation corrections, we use a versioning scheme inspired by https://semver.org/. Between each version only the data that has actually changed is updated, whereas unchanged data is referred to by links to older versions.

### MultiModal Browser

The MultiModal Browser (MMB) is written in Java and thereby executable on all major platforms (Linux, Mac, Windows). The code is publicly available in a GitHub repository: https://github.com/platybrowser/mmb-fiji. The the repository also contains the latest installation and usage instructions: https://github.com/platybrowser/mmb-fiji#platybrowser-fiji. For easy installation, we deploy the MMB as a Fiji plugin, which can be readily downloaded via the Fiji update site mechanism (Schindelin et al., 2012). The graphical user interface of the MMB is implemented using the Java Swing library (https://en.wikipedia.org/wiki/Swing_(Java)). The image data handling and visualisation is based on imglib2 (Pietzsch et al., 2012), a java library for processing large N-dimensional image data. For image data visualisation, the MMB uses imglib2’s default image viewer BigDataViewer (Pietzsch et al., 2015). For remote streaming of large image data, we employ n5-aws-s3, a flavour of the n5 file format (Pisarev and Saalfeld, 2020) using the amazon S3 object store API (https://en.wikipedia.org/wiki/Amazon_S3). At the moment of publication, n5-aws-s3 was not yet supported by the imglib2 and BigDataViewer eco-system as hosted on https://github.com/imglib and https://github.com/bigdataviewer. Thus, this functionality is currently provided by code within our own development branch: https://github.com/tischi/bigdataviewer-core/tree/n5-aws-s3. We however expect this functionality to be soon available within the official BigDataViewer repositor https://github.com/bigdataviewer/bigdataviewer-core. Also interactive visualisation of label-mask (segmentation) images and associated numeric tabular data as needed for the MMB was not available within the imglib2 eco-system and we devised our own solutions. Much of the relevant code was conceptualised and developed during a “Fiji hackathon” in Ostrava (Czech Republic, January 2019), and we are very grateful to Tobias Pietzsch, who, during this event, provided invaluable support with the software design. The resulting code for interactive visualisation of segmentation images and associated tabular data is currently hosted here: https://github.com/tischi/table-utils. We however already took first steps towards integrating these functionalities into the ImageJ eco-system. We would like to note that the above information is only relevant for software developers, while for end-users all necessary dependencies will be shipped automatically with the Fiji update site that is hosting the MMB.

### PlatyBrowser

The PlatyBrowser uses the MMB framework for the interactive browsing of our *Platynereis* atlas data. To this end, the MMB fetches data from the https://github.com/platybrowser/platybrowser-backend GitHub repository (see above section on Data Provision). In order to use the MMB for other data-sets, the layout in https://github.com/platybrowser/platybrowser-backend/tree/master/data needs to be replicated. The repository contains different versions of the data; the most recent version at time of publication is 1.0.0 and the reported results are based on this version unless noted otherwise.

