## Supplementary material for "Whole-body integration of gene expression and single-cell morphology"

### Supplementary Figures

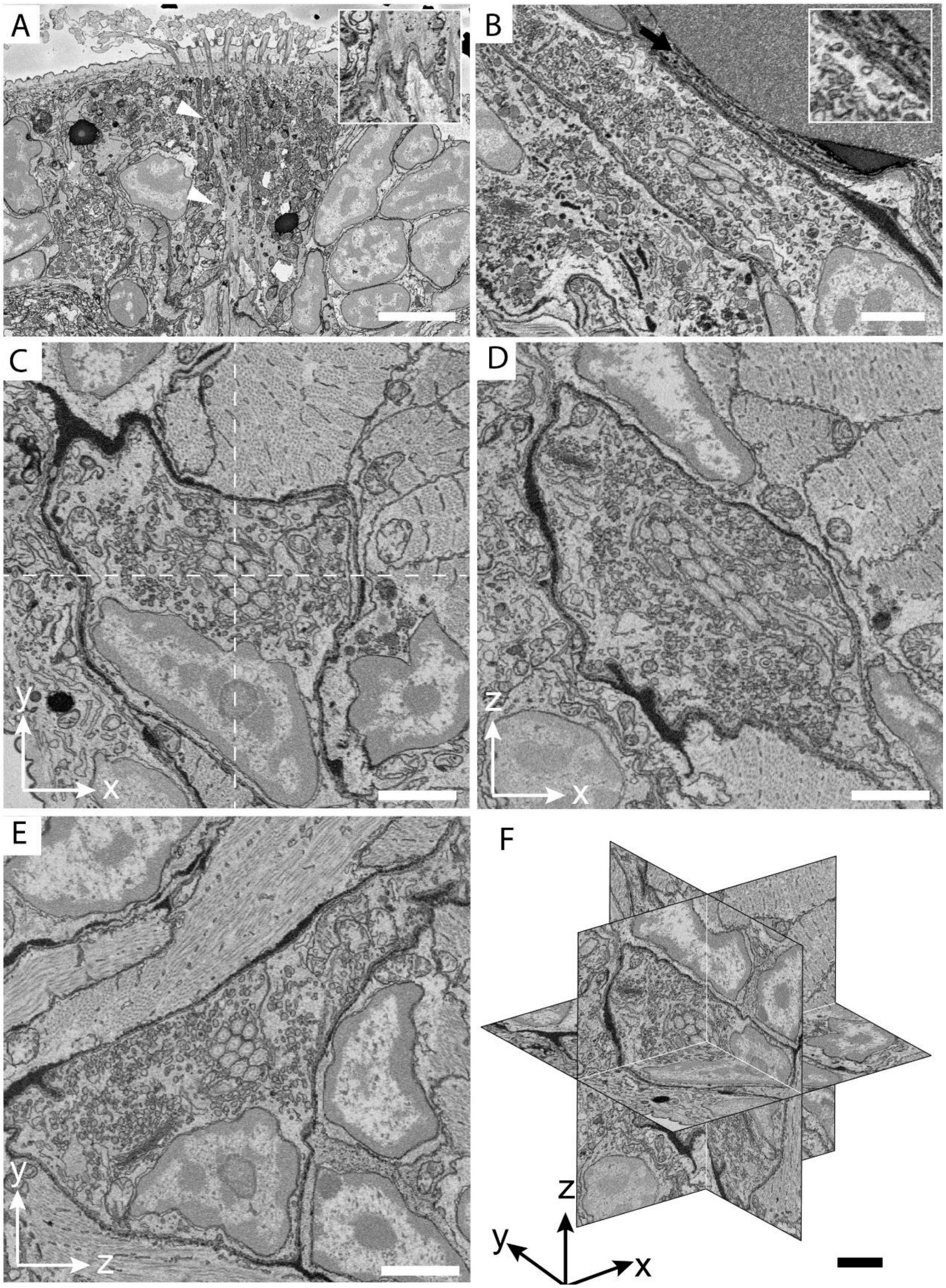

**Supplementary Figure 1. Ultrastructure of different cell types can be identified in the SBEM volume at native resolution.**

**A.** Ciliated support cells which are part of the nuchal organ in *Platynereis*. Cytoskeletal fibres (arrowheads) and anchoring points at the junction of the support cell and the underlying muscles are visible (inset) (Scale bar 5 µm). **B.** Cell of the nephridia which contains a lumen occupied by motile cilia. These cells contain numerous vesicles responsible for various forms of cell transport. A site of endocytosis, identified by the presence of a clathrin-coated pit is highlighted (arrow)(inset) (Scale bar 2 µm). **C-F.** Orthogonal projections of the image displayed in Figure 1E - scaling factor of 2.5x was applied to the Z plane to get an isotropic render. Panels A and B are snapshots selected from the full volume and can be retrieved as bookmarks in the PlatyBrowser. (Scale bars 2 µm).

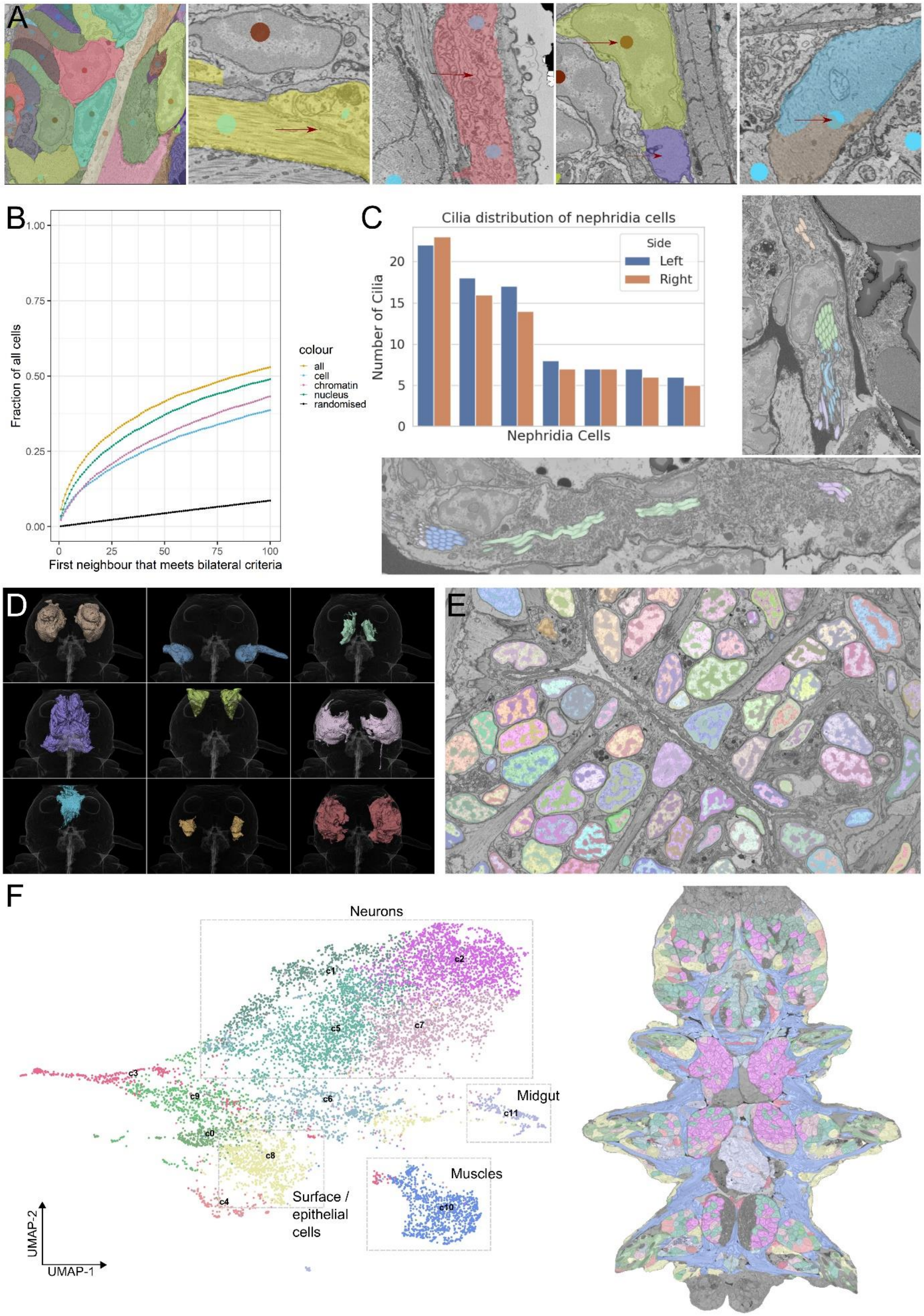

A
B
Fraction of all cells
First neighbour that meets bilateral criteria
colour
all
cell
chromatin
nucleus
randomised
C
Cilia distribution of nephridia cells
Side
Left
Right
Number of Cilia
Nephridia Cells
D
E
F
Neurons
Midgut
Muscles
Surface / epithelial cells
UMAP-2
UMAP-1
c0
c1
c2
c3
c4
c5
c6
c7
c8
c9
c10
c11

**Supplementary Figure 2. Segmentation validation; ultrastructure segmentation and morphological clustering**

**A:** Nuclei and cells were annotated by domain experts for 8 slices (4 transversal, 4 horizontal), see leftmost image for example annotations. We used these annotations to find false merge errors, see the two middle images with arrows highlighting the cell membrane not picked up, and false split errors, see two rightmost images with arrows highlighting parts of the cell that were split off, in the automated segmentation. **B:** Bilateral pairs analysis - the graph shows the fraction of all cells that find a potential bilateral partner within a certain number of neighbours. ‘Neighbours’ being all cells ranked by their distance in morphology space from 1 = very closest in morphology space to 10345 = the very furthest away in morphology space. E.g. an x,y position of 20, 0.5 would mean that 50% of cells find a potential bilateral partner within 20 neighbours. ‘All’ uses all morphology features in **Supplementary Table 2**, ‘cell’ uses only the cell features, ‘nucleus’ uses all nucleus and chromatin features and ‘chromatin’ uses only intensity and texture features of chromatin. The randomised line was generated by random shuffling of the ranking of cells in morphology space, and taking the mean of 100 trials. **C:** The distribution of cilia per cell for the nephridia on both sides is stereotypical as can be seen from the plot. Cilia in a given cross-section of the lumen almost exclusively start off from the same cell, see segmented cilia coloured by their cell of origin overlaid on the EM; upper image shows a cross section of the right nephridium, lower image of the left nephridium. **D:** 3D rendering of all segmented ganglia in the head of *Platynereis* (colours match upper row of Figure 5D). **E:** Chromatin segmentation overlaid on the EM. The dark phase (classical heterochromatin and nucleolus) and the light phase (classical euchromatin) are segmented. **F:** Left - UMAP of all cells based on all morphological features. Points are coloured by membership to different morphological clusters (c0-c11). Grey rectangles highlight some clusters of interest that correspond to certain regions / tissue types. Right - morphological clusters overlain on a section of the EM dataset. For panel E and F (right), bookmarks are available in the PlatyBrowser.

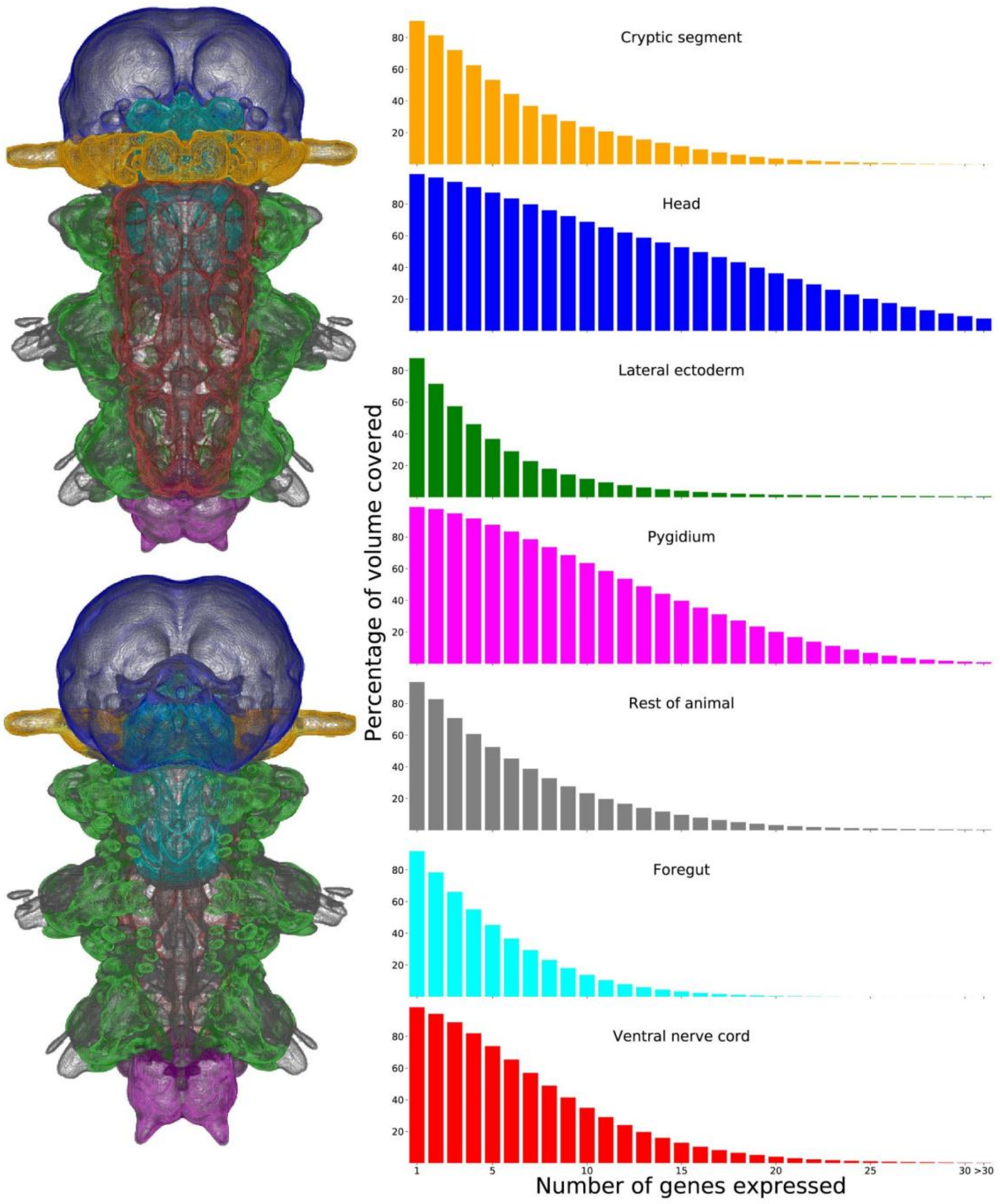


**Supplementary Figure 3. Coverage of gene expression by region in the ProSPr atlas.**

Quantification of gene coverage by animal region. Animal regions are coloured as indicated in the 3D views. Histograms represent the percentage of volume containing signal for the number of genes indicated in the x axis.

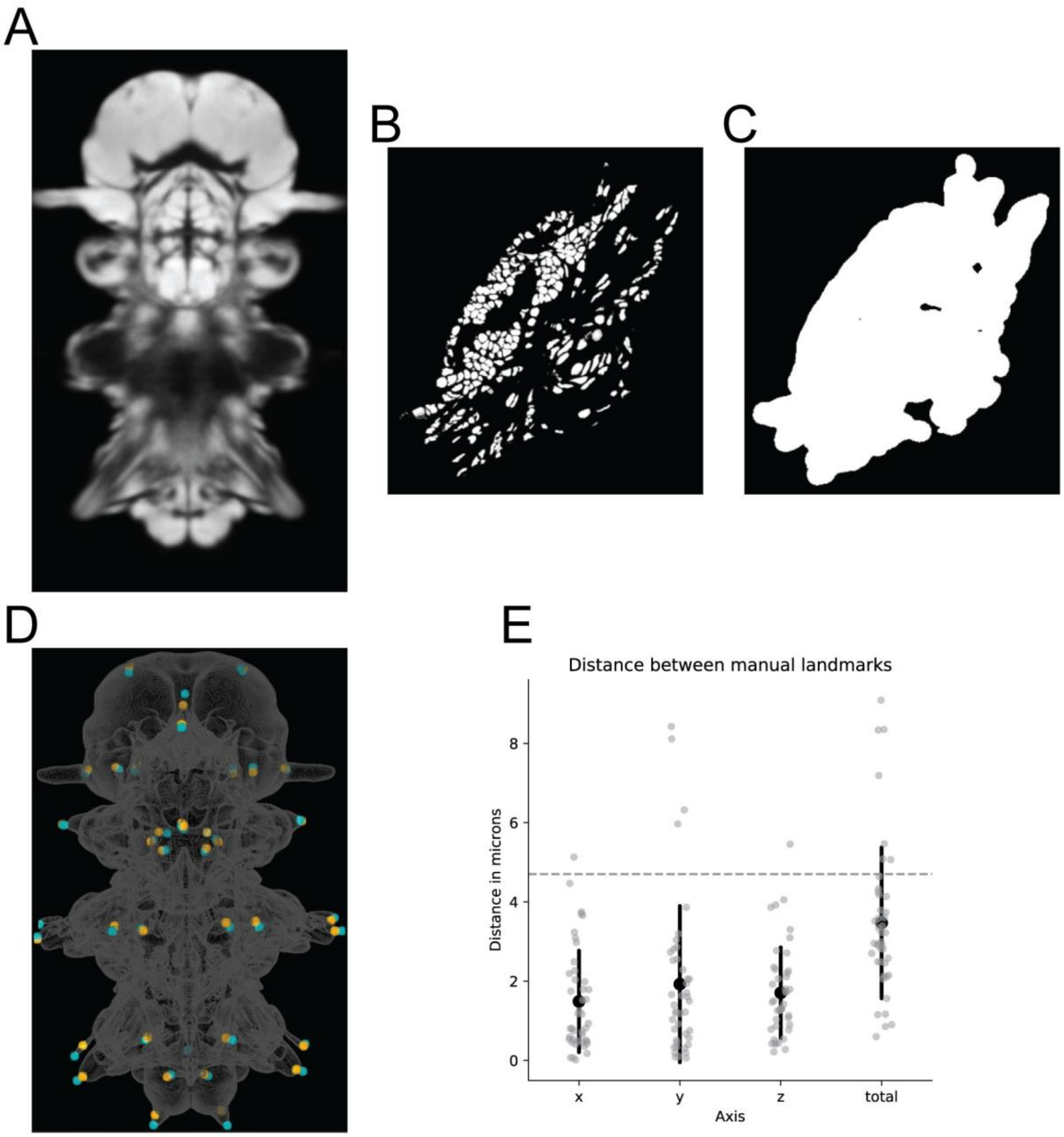


**Supplementary Figure 4. Registration of ProSPr average DAPI signal to segmented EM nuclei.**

**A-C**: Exemplary single planes of the image data stacks, which were used as input to the registration. **A**: DAPI: Average DAPI signal of 153 images from the ProSPr atlas. **B:** EM-Nuclei: Mask of segmented nuclei of the EM individual. **C**: EM-Mask: Binary mask, created by dilation and binarisation of the EM-Nuclei image. The EM-Mask was used to restrict the elastix optimisation algorithm to relevant parts of the image. **D**: three-dimensional visualisation of the overlay after final registration (see methods) between the 43 manually selected landmarks in both datasets. Landmarks in the SBEM dataset are plotted in orange, and landmarks in the ProSPr atlas are plotted in cyan. Spheres have a diameter of 5 microns. Plotted in ProSPr space. The gray outline is an arbitrary mask extracted from the DAPI signal. **E**: quantification of the distance between the 43 landmarks in each axis (ProSPr atlas space: 'x' corresponds to medio-lateral, 'y' to anterior-posterior, and 'z' to dorso-ventral axis), and in total. Horizontal dashed line represents the average cell diameter.

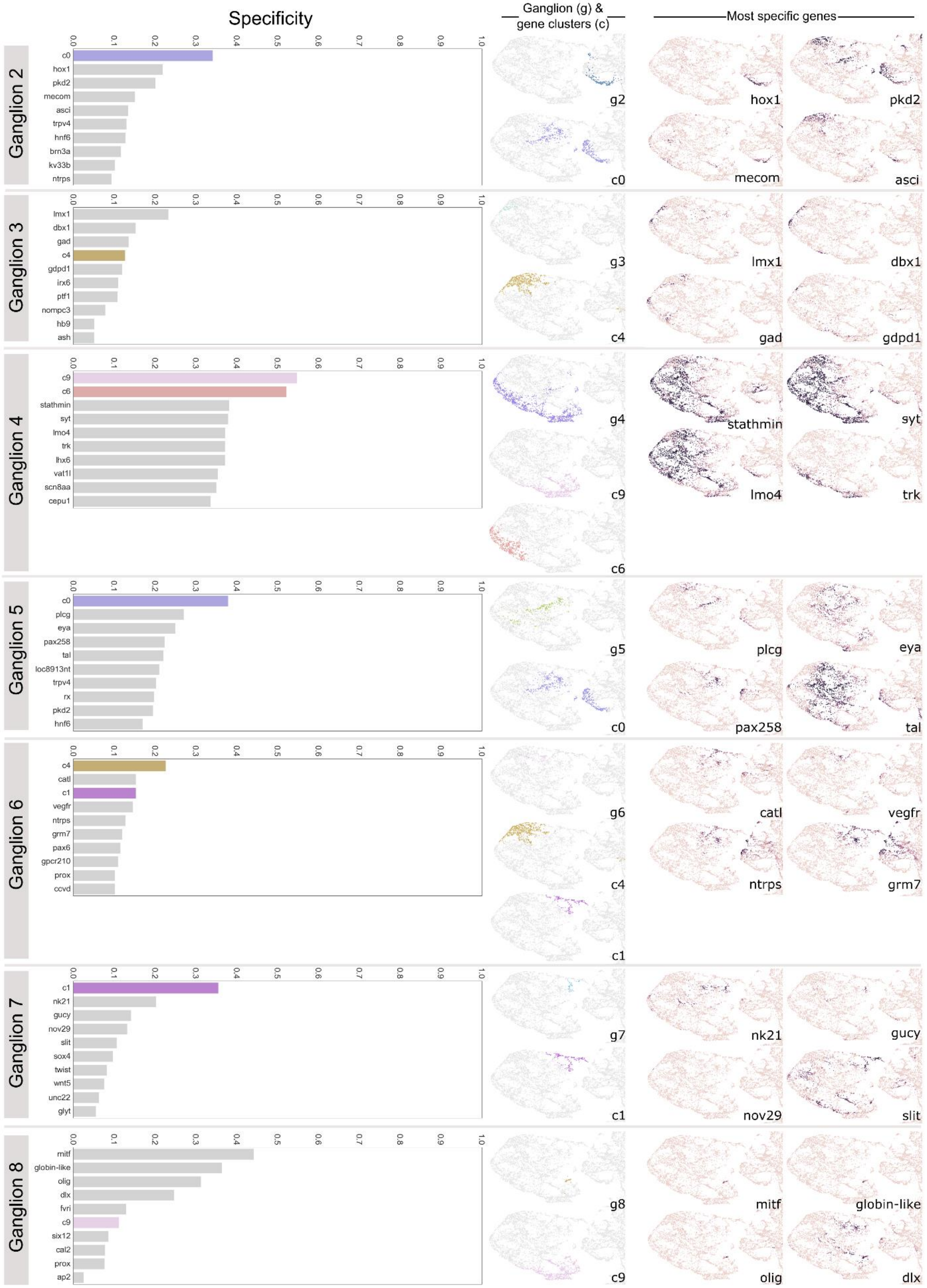

Specificity
Ganglion (g) & gene clusters (c)
Most specific genes
Ganglion 2
c0 hox1 pkd2 mecom asci trpv4 hnf6 brn3a kv33b ntrps
g2 c0 hox1 pkd2 mecom asci
Ganglion 3
lmx1 dbx1 gad c4 gdpd1 irx6 ptf1 nompc3 hb9 ash
g3 c4 lmx1 dbx1 gad gdpd1
Ganglion 4
c9 c6 stathmin syt lmo4 trk lhx6 vat1l scn8aa cepu1
g4 c9 c6 stathmin syt lmo4 trk
Ganglion 5
c0 plcg eya pax258 tal loc8913nt trpv4 rx pkd2 hnf6
g5 c0 plcg eya pax258 tal
Ganglion 6
c4 catl c1 vegfr ntrps grm7 pax6 gpcr210 prox ccvd
g6 c4 c1 catl vegfr ntrps grm7
Ganglion 7
c1 nk21 gucy nov29 slit sox4 twist wnt5 unc22 glyt
g7 c1 nk21 gucy nov29 slit
Ganglion 8
mitf globin-like olig dlx fvri c9 six12 cal2 prox ap2
g8 c9 mitf globin-like olig dlx
0.0 0.1 0.2 0.3 0.4 0.5 0.6 0.7 0.8 0.9 1.0

**Supplementary Figure 5. Specificity of gene clusters and individual genes for head ganglia.**

Comparison of specificity of gene clusters and individual genes for remaining ganglia not covered in figure 5 (g2-g8). Left column – graphs of top 10 scoring genes (grey bars) or gene clusters (coloured bars) by F1 specificity score (see methods). Right – zooms of the head region of the UMAP from figure 5A coloured by ganglia (g), top scoring genetic clusters (c), or gene expression overlap value (0-1, same scale as in figure 5E) for top scoring genes. Note: here we show only the head region of the UMAPs, for easier comparison, but some genes and gene clusters have expression domains outside of the head which contribute to their lower specificity scores.

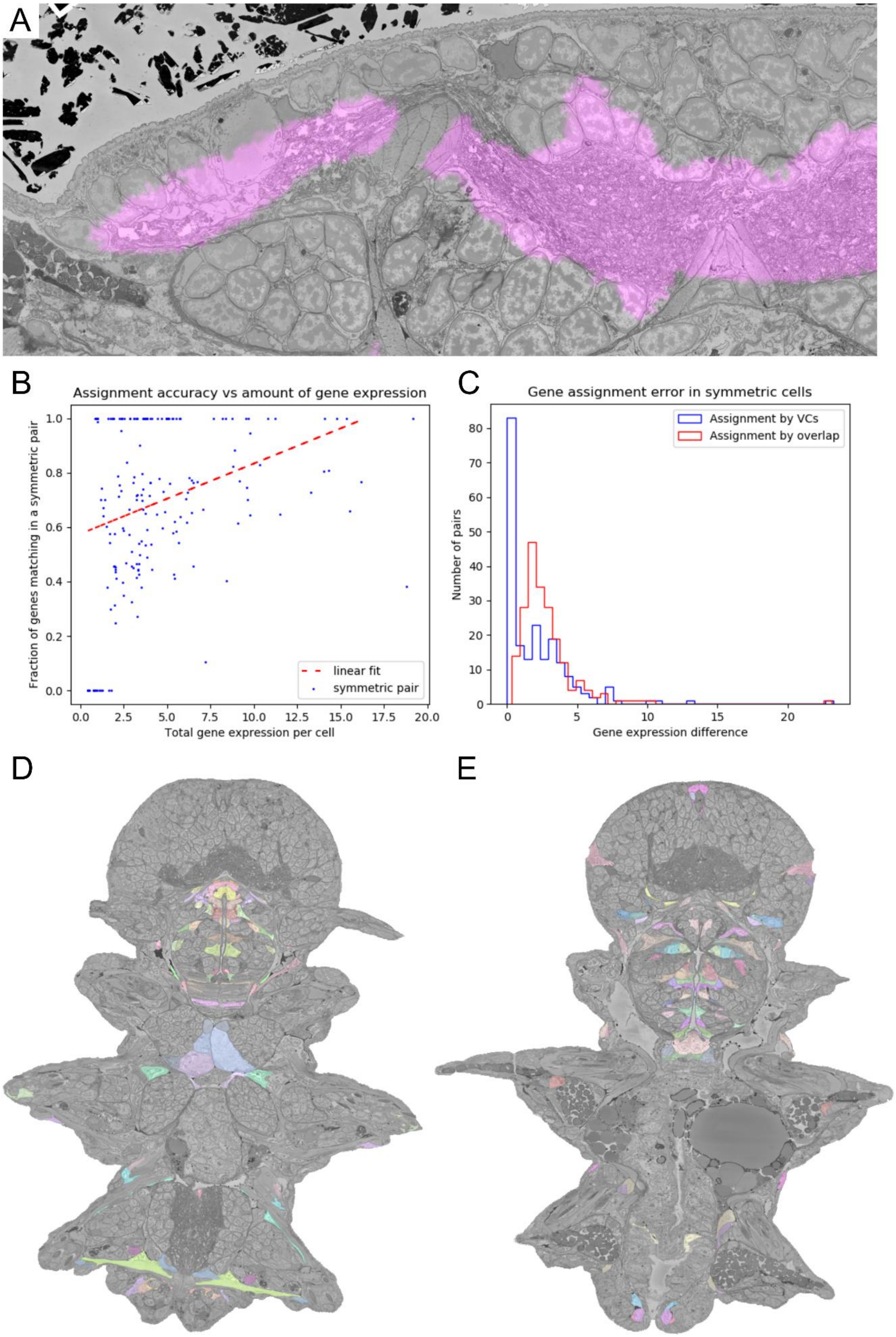
A
B
Assignment accuracy vs amount of gene expression
Fraction of genes matching in a symmetric pair
Total gene expression per cell
linear fit
symmetric pair
C
Gene assignment error in symmetric cells
Number of pairs
Gene expression difference
Assignment by VCs
Assignment by overlap
D
E

**Supplementary Figure 6. Gene expression assignment**

**A**: 'Gene leaking' of *glt1* (glutamate receptor). While the true expression is confined to the neuropil, the bordering regions such as neural somas, muscles and epithelial cells also show a high level of expression, originating from sample variability and limited registration accuracy. **B:** Dependency of the assignment accuracy on the total level of gene expression in a cell (the fractions of gene expression for each gene summed up). The assignment performed better for the cells in gene rich areas. **C**: Assignment errors in the symmetric cells pairs for the Virtual Cells assignment and assignment by overlap. Assignment error is defined as the absolute difference in gene expressions assigned to the cells of a symmetric pair. **D-E**: Examples of symmetric cells pairs – cells with similar mirror location and morphology, supposedly representing the same cell type and expressing the same genes.

### Supplementary Tables

| Region or tissue name | Number of cells | Type |
|---|---|---|
| Cryptic segment | 430 (3.8 %) | Region |
| Foregut | 1458 (12.8 %) | Region |
| Glands | 87 (0.8 %) | Tissue |
| Head | 3754 (32.9 %) | Region |
| Lateral ectoderm | 1510 (13.2 %) | Region |
| Midgut | 304 (2.7 %) | Region |
| Muscle | 988 (8.7 %) | Tissue |
| Pygidium | 387 (3.4 %) | Region |
| Ventral nerve cord | 2111 (18.5 %) | Region |

**Supplementary Table 1: Number of cells per region/tissue**

The number of (segmented) cells per region / identifiable tissue type. Note that the region delineations are derived from ProSPr and do not align perfectly with the EM.

| Feature name | Segmentations | Description |
|---|---|---|
| shape_volume_in_microns | Cell, nucleus, chromatin | Volume in cubic microns |
| shape_extent | Cell, nucleus, chromatin | Ratio of pixels in the object to pixels in the total bounding box |
| shape_equiv_diameter | Cell, nucleus, chromatin | Equivalent diameter - the diameter of a sphere with the same volume as the object |
| shape_major_axis | Cell, nucleus, chromatin | Length of the major axis of the fitted ellipsoid |
| shape_minor_axis | Cell, nucleus, chromatin | Length of the minor axis of the fitted ellipsoid |
| shape_surface_area | Cell, nucleus, chromatin | Surface area of object mesh (mesh calculated by the Lewiner marching cubes algorithm) |
| shape_sphericity | Cell, nucleus, chromatin | Measure of how spherical an object is (0-1 scale, with 1 being a perfect sphere) calculated as $36\pi V^2/S^3$ where V is the volume of the object, and S is its surface area. |
| shape_max_radius | Cell, nucleus, chromatin | Maximum distance from a pixel within the object to the outside (Euclidean distance) |
| intensity_mean | Cell, nucleus, | Mean intensity of the segmented object |

| | | |
|---|---|---|
| | chromatin | |
| intensity_st_dev | Cell, nucleus, chromatin | Standard deviation of intensity of the segmented object |
| intensity_median | Cell, nucleus, chromatin | Median intensity of the segmented object |
| intensity_iqr | Cell, nucleus, chromatin | Interquartile range (iqr) of intensity of the segmented object |
| intensity_total | Cell, nucleus, chromatin | Sum of intensity values of the segmented object |
| Intensity_mean_ (25/50/75/100) | Nucleus | Mean intensity of different radial zones of the segmented object. These zones are calculated from a euclidean distance transform of the object, that is normalised to run from 0 to 1 (1 being the point in the object furthest from the edge). Zone 25 is then the outermost 25% (values 0-0.25), zone 50 from 25% to 50% (values 0.25-0.5), zone 75 from 50% to 75% (values 0.5-0.75) and zone 100 from 75% to 100% (0.75-1.0) |
| Intensity_st_dev_ (25/50/75/100) | Nucleus | Standard deviation of intensity of different radial zones of the segmented object (see intensity_mean_(25/50/75/100) for how these zones are calculated) |
| Intensity_median_ (25/50/75/100) | Nucleus | Median intensity of radial zones of the segmented object (see intensity_mean_(25/50/75/100) for how these zones are calculated) |
| Intensity_iqr_ (25/50/75/100) | Nucleus | Interquartile range (iqr) of intensity of different radial zones of the segmented object (see intensity_mean_(25/50/75/100) for how these zones are calculated) |
| Intensity_total_ (25/50/75/100) | Nucleus | Sum of intensity values of different radial zones of the segmented object (see intensity_mean_(25/50/75/100) for how these zones are calculated) |
| texture_hara(1-13) | Cell, nucleus, chromatin | Haralick texture features of the segmented object. Haralick texture features 1-13 are commonly used texture descriptors in image analysis - each is a statistic derived from the grey level co-occurrence matrix of an image. (Haralick et al., 1973) |
| shape_edt_mean_(het/eu) _nucleus | Chromatin | Mean of values in normalised euclidean distance transform (edt) of whole nucleus covered by the current segmented object (either euchromatin or heterochromatin + nucleolus segmentation). This is a measure of the distribution of chromatin within the nucleus (low values indicate a distribution mostly towards the edge of the nucleus, while higher values indicate a distribution closer to the centre). The euclidean distance transform is calculated for the whole nucleus segmentation, and normalised to run from 0 to 1 (1 being the point furthest from the edge). |
| shape_edt_stdev_(het/eu) _nucleus | Chromatin | Standard deviation of values in normalised euclidean distance transform (edt) of whole nucleus covered by the current segmented object (either euchromatin or heterochromatin + nucleolus segmentation). This is a measure of the distribution of chromatin within the nucleus (high values indicate a varied distribution in the nucleus i.e. some towards the outside of the nucleus, as well as some towards the centre). The euclidean distance transform is |

| | | |
|---|---|---|
| | | calculated for the whole nucleus segmentation, and normalised to run from 0 to 1 (1 being the point furthest from the edge). |
| Shape_edt_median_(het/eu)_nucleus | Chromatin | Same as shape_edt_mean_(het/eu)_nucleus, but using median rather than the mean |
| shape_edt_iqr_(het/eu)_nucleus | Chromatin | Same as shape_edt_stdev_(het/eu)_nucleus, but using interquartile range (iqr) rather than standard deviation. |
| Shape_percent_(25/50/75/100)_(het/eu)_nucleus | Chromatin | Percent of different radial zones in the nucleus that are filled by either the heterochromatin+nucleolus or euchromatin segmentation - this is a measure of the distribution of chromatin within the nucleus. The radial zones are calculated from a euclidean distance transform of the whole nucleus, that is normalised to run from 0 to 1 (1 being the point furthest from the edge). Zone 25 is then the outermost 25% (values 0-0.25), zone 50 from 25% to 50% (values 0.25-0.5), zone 75 from 50% to 75% (0.5-0.75) and zone 100 from 75% to 100% (0.75-1.0) |

**Supplementary Table 2: Description of morphological features used in clustering analysis**

Description of morphological, intensity and texture features used in clustering analysis. Some features are calculated for a number of different segmentations (as indicated by the second column). Het / eu in feature names are short for 'heterochromatin + nucleolus' and 'euchromatin' respectively. The full table of calculated features for cells is available here: https://github.com/platybrowser/platybrowser-backend/blob/master/data/1.0.0/tables/sbem-6dpf-1-whole-segmented-cells/morphology.csv and for nuclei (+ chromatin) here: https://github.com/platybrowser/platybrowser-backend/blob/master/data/1.0.0/tables/sbem-6dpf-1-whole-segmented-nuclei/morphology.csv. The clustering derived from both these sets of features combined is available here: https://github.com/platybrowser/platybrowser-backend/blob/master/data/1.0.0/tables/sbem-6dpf-1-whole-segmented-cells/morphology_clusters.csv
